# Low accuracy of complex admixture graph inference from *f*-statistics

**DOI:** 10.1101/2025.03.07.642126

**Authors:** Lauren E. Frankel, Cécile Ané

## Abstract

*F*-statistics are commonly used to assess hybridization, admixture or introgression between populations or deeper evolutionary lineages. Using simulations, we find that network complexity had a large impact on the accuracy to infer the network structure from *f* statistics. Networks recovered accurately had one reticulation, or had their reticulations in “large” cycles of at least 4 nodes in all subnetworks. But accuracy was extremely poor to infer complex networks, in which a reticulation is part of a small cycle of only 3 nodes in some subnetwork. Accuracy also decreased with increasing number of reticulations and the network level. For these networks, accuracy was low even from large data sets with low mutation rate, under a molecular clock, and retaining many top-scoring graphs. Yet in all cases, the network’s major tree was recovered reliably. When the molecular clock was violated, the *f*_4_-test tended to falsely detect the presence of reticulation in large data sets or under a high mutation rate. Rate variation also impacted network inference accuracy and increased the rate of falsely rejecting 1 reticulation as being adequate. We propose that identifiability, or lack thereof, is underlying the contrasting recoverability between simple and complex networks. Our findings suggest that the major tree is one feature that might be estimable from *f*-statistics. In practice, we recommend evaluating a large set of top-scoring networks inferred from *f*-statistics, and even so, using caution in assuming that the true network is part of this set. The extent of rate variation should be assessed in the system under study, especially at deeper time scales, or when using fast-evolving loci.

## 1 Introduction

The *f*_4_ statistic is very popular to identify histories of introgression or admixture and is the basis for many high-profile studies (Fig. 1). It uses allele frequencies from 4 populations and measures the correlation of allele frequency differences between pairs of populations [Peter, 2016, Soraggi and Wiuf, 2019, Lipson, 2020, Peter, 2022]. The *D*-statistic, also widely used, is a normalized version of *f*_4_ [Patterson et al., 2012, Green et al., 2010]. *F*-statistics form the basis of many discoveries about archaic human population dynamics [Lipson et al., 2018, Wang et al., 2021, Flegontov et al., 2019], especially as ancient DNA extraction and sequencing techniques have advanced in recent years. *F*-statistics have also been used to study a variety of other organisms (for example, in domestication of horses, Librado et al. [2021]; mammoths, van der Valk et al. [2021]; wild grapes, Zecca et al. [2020]). From *f*_4_ statistics, admixture graphs can be readily and rapidly fitted to data and inferred, using tools such as qpgraph [Patterson et al., 2012], qpWave [Haak et al., 2015], qpAdm [Haak et al., 2015, Harney et al., 2021] and `find_graphs` [Maier et al., 2023] all implemented in the R package admixtools [Patterson et al., 2012, Maier et al., 2023], as well as poolfstat [Gautier et al., 2022] and AdmixtureBayes [Nielsen et al., 2023].

**Figure 1:**
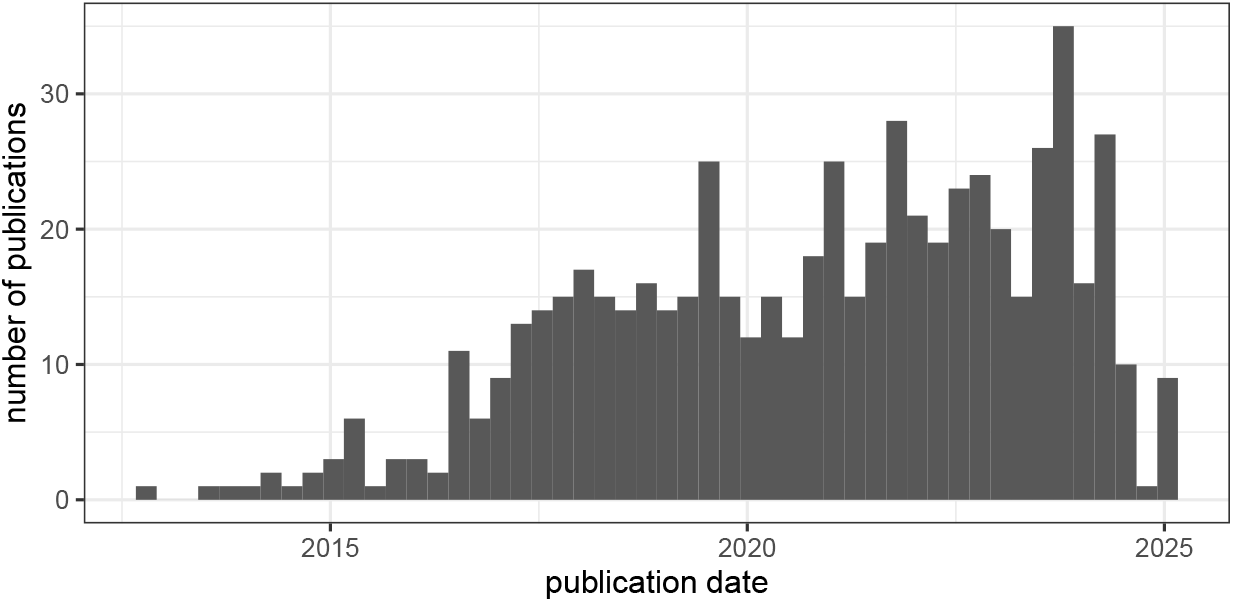
Number of publications in the OpenAlex catalog [Priem et al., 2022] with full text including ‘admixtools’, ‘admixtools2’ or ‘f4 statistics’. Of these 616 publications, 110 were published in *Science* or in one of the following Nature journals: *Scientific Reports, Nature, Nature Communications, Nature Genetics, Nature Ecology & Evolution* and garnered a total of 14,770 citations. Data were retrieved using the R package openalexR [Massimo et al., 2024] on 2025-02-08.

In view of the many high-profile studies featuring admixture graphs inferred from *f*-statistics, surprisingly few studies have evaluated the accuracy of admixture graph inference. In simulations, Maier et al. [2023] found the recovery of the true graph as the best-scoring graph to decline dramatically as the number of admixture events increases, or as the number of populations decreases. In our work, we evaluate other factors that may contribute to accuracy of graph inference, including admixture proportions and various aspects of graph complexity beyond the number of introgression events, such as the network level and the size of cycles associated with each event. More *complex* networks are networks with more reticulations, of higher level, and with one or more reticulations that are part of small cycles of only 3 nodes when other reticulations are removed (explained below). We consider *simpler* networks as having few reticulations, separated from each other, and that induce cycles of 4 or more nodes.

One underlying assumption of the *f*_4_ test of admixture, and of other hybridization tests (for example, the *D*-statistic, Durand et al. [2011], Patterson et al. [2012]; *D*_3_, Hahn and Hibbins [2019]; HyDe, Blischak et al. [2018]), is a constant substitution rate across genes and/or across lineages. Lineage-specific substitution rate variation occurs when genome-wide rates differ between population lineages, due to shifts in physiological or life-history traits between populations or other factors that affect the generation time or the per-generation mutation rate [Lehtonen and Lanfear, 2014], both of which have been documented to vary in diverse groups of organisms, even between closely related lineages [Smith and Donoghue, 2008, Thomas et al., 2010, Wang and Obbard, 2023]. When these tests are applied at a macroevolutionary scale where substitution rates are likely to vary, this could present a problem in the accuracy and interpretation of admixture detection and admixture histories. Even at short evolutionary timescales substitution rates can vary due to variation in effective population size combined with selection, or in traits such as generation interval [Petit and Barbadilla, 2009, Abu-Elmakarem et al., 2024, Gossmann et al., 2012], which have been documented to vary over different timescales between archaic and modern humans [Wang et al., 2023, Coll Macià et al., 2021, Ragsdale et al., 2023b].

In simulations, Frankel and Ané [2023] found that substitution rate variation across lineages (rather than across genes or lineage-by-genes) can be problematic for several methods based on 3-taxon and 4-taxon summary statistics. When certain lineages evolve faster or slower than others on average (across the genome), then hybridization summary statistics, including the *D*-statistic, often erroneously indicate there is hybridization when there is none. Frankel and Ané [2023] also found that admixture graph complexity poses a problem to detect hybridization with summary statistics. Specifically, having 2 or more hybridization events in a 10-taxon graph lowered power, even in the absence of rate variation. If *f*_4_ behaves similarly to *D*, evolutionary histories with 2 or more hybridization events may not be accurately estimated by qpgraph or `find_graphs`.

In this work, we present an extensive simulation study to quantify the performance of *f*_4_-based admixture graph inference using `find_graphs`. We identify specific measures of network complexity that strongly impact the inference accuracy of the graph topology. We also assess how violation of the molecular clock affects the *f*_4_-test and the *f*_4_ worst residuals used for selecting an appropriate number of reticulations.

In this study, to simplify terminology, we use ‘graph’ and ‘network’ interchangeably, and we denote by ‘reticulation’ any biological process that results in a populations inheriting genetic material from multiple parent populations, including admixture, gene flow, and hybridization. We also denote a node with multiple parent populations (e.g. an admixed population) as a ‘hybrid’ node, regardless of the biological process. Similarly, edges leading to a hybrid node are call hybrid edges, also known as admixture edges when the process is known to be population admixture.

## 2 Methods

### 2.1 Simulation

#### 2.1.1 Networks and simulation parameters

Under a large number of parameter combinations, we simulated data along one of two fixed “true” networks, chosen for being realistic networks of relevance to empirical data (Fig. 2). The more complex network came from Flegontov et al. [2023] Fig. 3A, with edge parameters (number of generations, population size, admixture weights) detailed in in their S13 Table. This network includes a chimpanzee, Denisovan, two Neanderthal, two African, and two non-African populations. It also includes a bottleneck event and four hybridization events, so we call it `g4` here. To assess the effect of network complexity on inference, we modified this `network g4` by pruning the 3 minor hybrid edges with lowest admixture weights (*γ* = 0.06, 0.19 and 0.32 in orange in Fig. 2). This left just one reticulation event (with minor *γ* = 0.48, in blue) ancestral to the Denisovan population – so we call this network `g1`.

**Figure 2:**
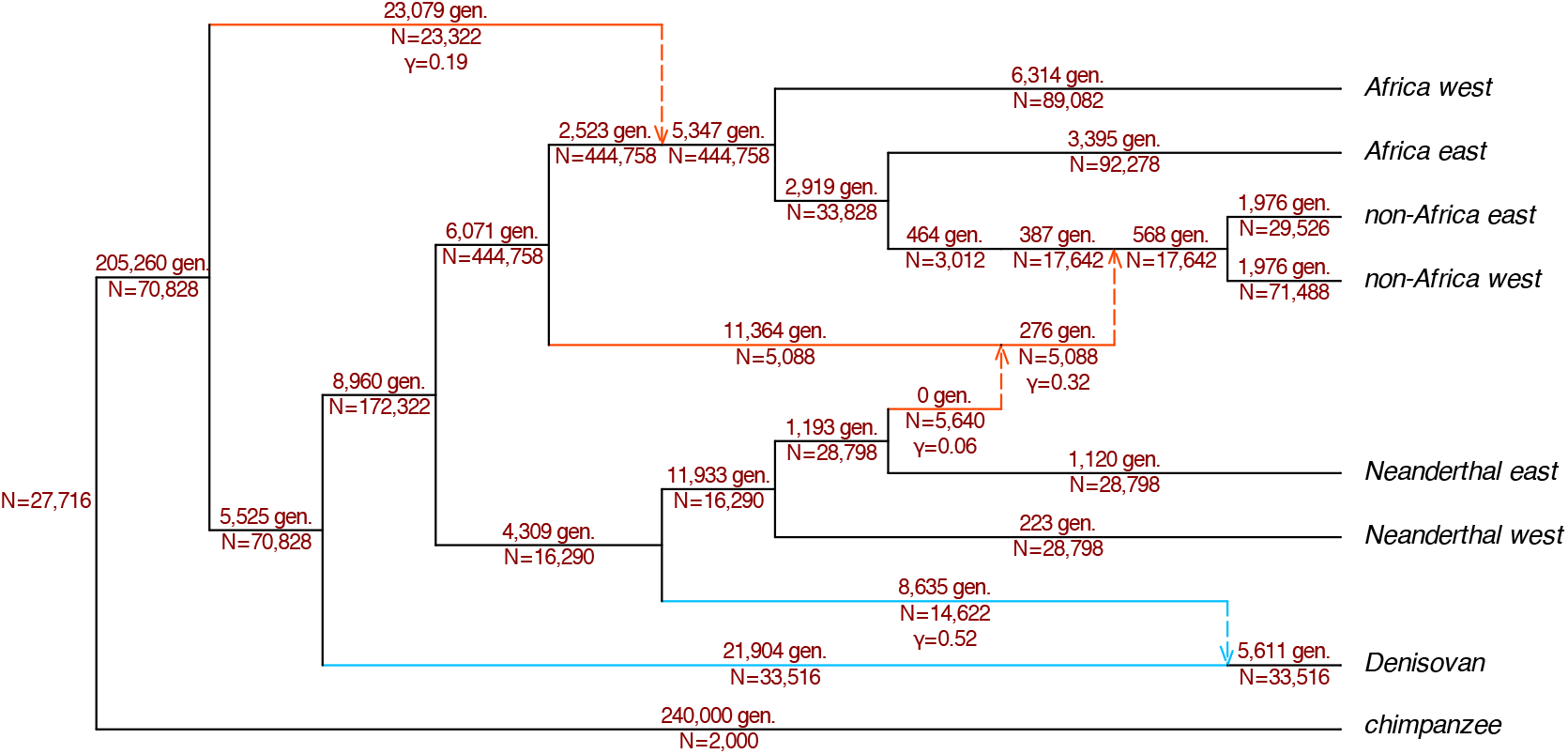
Network `g4` used in simulations, with *h* = 4 reticulations [from Flegontov et al., 2023, see Methods]. Edges are annotated with number of generations, effective haploid population size *N* and inheritance probability *γ*, also called admixture weight, for hybrid edges. The network `g1`, also used under many parameter combinations, is the subnetwork of `g4` obtained by keeping *h* = 1 reticulation of large effect (inheritance *γ* = 0.48 and 0.52 from its parent populations, edges shown in blue), and removing the 3 reticulations of lesser effect (which removes edges shown in orange). See Fig. S1 for additional networks used to simulate data under baseline parameters.

**Figure 3:**
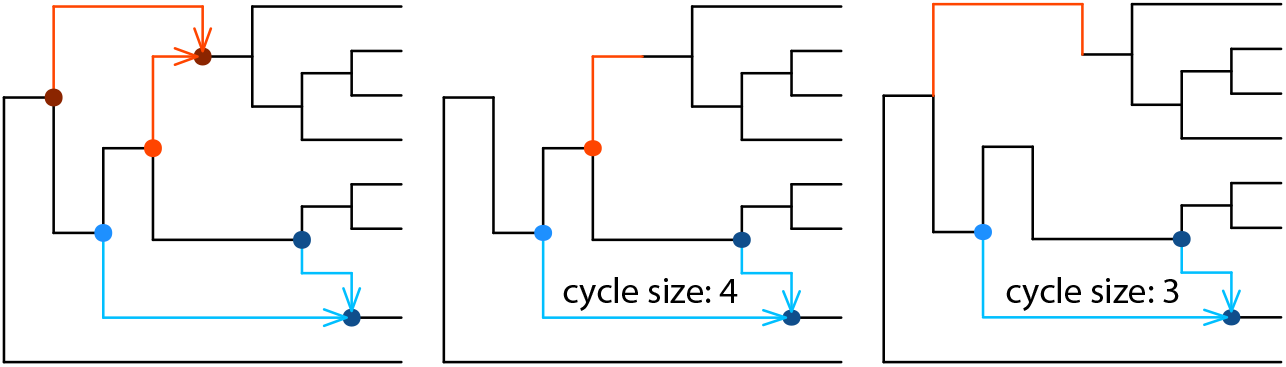
Left: one of the networks used to simulate data. It has *h* = 2 reticulations in one blob, whose nodes are marked by circles, so its level is *ℓ* = 2. This network is tree-child and galled. Middle: subnetwork obtained by removing one of the parental edge to the red reticulation. Only the blue reticulation remains, in a cycle of 4 nodes (and 4 edges). Right: subnetwork obtained by removing the other red parental edge, leaving the blue reticulation in a cycle of 3 nodes. The minimum cycle size of the blue reticulation is then *c* = 3. The red reticulation also has *c* = 3 (subnetworks not shown).

For one baseline combination of parameter values, we simulated data under 16 additional population networks, with varying levels of complexity (Fig. S1). First, to assess the effect of parental inheritance probabilities *γ*, we decreased the minor *γ* on `g1` from 0.48 to 0.06, to match the smallest *γ* on `g4`. Accordingly, we increased the two smallest *γ*s on `g4` from 0.06 and 0.19 both to 0.32. The other 14 networks all have large inheritance probabilities (*γ* ≥ 0.32) but other topologies with 1 to 4 reticulation events, and various complexity features such as level and cycle sizes, described next. They were all transformed from `g4`, with similar edge parameters (number of generations and population size).

#### 2.1.2 Measures of network complexity

Levels and cycle sizes are features are known to affect network identifiability from various data types [e.g. Solís-Lemus and Ané, 2016, Baños, 2019, Xu and Ané, 2023, Allman et al., 2024, Englander et al., 2025, Allman et al., 2025]. These features are briefly described here, but more details can be found in the references above.

The *reticulation number h* of a network is the minimum number of edges that need to be deleted to obtain a tree. If each admixed population has exactly 2 parental populations, then this is simply the number of reticulation events. The *level ℓ* of a network is the maximum reticulation number within a *blob*. A *blob* is a maximal subgraph such that deleting any edge in it does not disconnect its nodes (considering edges as undirected). For example, `g4` in Fig. 2 has a single blob containing all *h* = 4 reticulations, so its level is *ℓ* = 4. In Fig. 3, the network on the left has a single blob (nodes marked by circles and edges connecting them). This blob contains its *h* = 2 reticulations, so its level is *ℓ* = 2. In all networks *ℓ* ≤ *h*, but we may have *ℓ < h* if some groups of reticulations are ‘separated’ from others (in cycles that do not share any edges), as in several networks used in simulations (Fig. S1). In a level-1 network in particular, reticulations are in cycles that do not overlap. Any network with 1 reticulation, like `g1`, is of level 1.

For each reticulation event *v*, we consider all the ways to isolate *v* by removing the other reticulations. There are 2 ways to remove a reticulation: by keeping either one of its parental populations, and deleting the edge representing inheritance from the other parental population. In each subnetwork in which only reticulation *v* remains, the subnetwork has a single cycle. The size of this cycle is its number of nodes (considering the network semidirected, in which the root is suppressed). The smallest cycle size, over all subnetworks in which only *v* remains, is called the *minimum cycle size* for *v*. For example, Fig. 3 (left) shows a network with 2 reticulations. If we consider the bottom reticulation *v*, shown with blue circles and blue parental lineages, *v* is part of 2 subnetworks, from the 2 ways to remove the other reticulation (top, in red). One of them has a cycle of size 4 (Fig. 3, center) and the other has a cycle of size 3 (Fig. 3, right), so *v* has minimum cycle size *c* = 3.

We count events with *c* ≤ 3, as this means low taxon sampling around the reticulation. Gene flow between sister species produce cycles of size *c* = 3 and are known to be either non-identifiable, or difficult to infer, depending on the data type. Identifiability can be helped if the network is *tree-child* (all nodes have at least one non-reticulate child) and *galled* (each reticulation is part of a unique cycle). In Fig. 3 (left), the red reticulation also has minimum cycle size *c* = 3, such that the network has 2 reticulations with *c* = 3. It is tree-child and galled, yet does not meet the cycle size requirement to be in the C_4_ class of Allman et al. [2025]. We will see that this network is recovered with low accuracy. In `g4` (Fig. 2), 2 reticulations have *c* = 3 (the reticulation in blue, and the reticulation with minor *γ* = 0.19) and the other 2 reticulations have *c* = 4. In `g1`, the reticulation has *c* = 4.

In networks used to simulate data (Fig. S1), the networks have various combinations of level (from 1 to 4) and number of reticulations with minimum cycle size *c* = 3 (from 0 to 2), to assess which of reticulation number, level, and minimum cycle size most affect network accuracy.

#### 2.1.3 Simulating gene trees

From `g1` and `g4`, we simulated sequence data for 1, 2, or 10 haploid individuals per population, and for various values of mutation rate and level of rate variation (see below) with either 11,800 or 118,000 gene trees. From all other networks, we simulated under the “baseline” best-case scenario, with 10 haploid individuals per population, a molecular clock, the slower mutation rate, and 118,000 gene trees.

For each combination of parameters, we simulated gene trees under the network multispecies coalescent (NMSC, Degnan 2018) using `PhyloCoalSimulations` v0.1.2 [Fogg et al., 2023, Fogg and Ané, 2022], written in Julia [Bezanson et al., 2017]. Simulating 11,800 or 118,000 gene trees allowed for a “buffer” of 1,800 or 18,000 gene trees respectively, that resulted in multiallelic sites (instead of biallelic) that cannot be used in *f*_4_ calculations. Gene trees had degree-2 nodes to mark each speciation and reticulation event, which allowed mapping each edge in a gene tree to the population lineage in the species network, that the gene edge evolved in.

#### 2.1.4 Simulating rate variation across population lineages

On `g1` and `g4`, we used five different levels of rate variation across population lineages. The distribution of rates across lineages was set to a log-normal distribution with mean 1 and one of 5 standard deviation *σ* on the log-scale: *σ* = 0 (a molecular clock), 0.15, 0.3, 0.5, or 0.7. The 0.1 and 0.9 quantiles of each distribution are (0.82, 1.2) for *σ* = 0.15; (0.65, 1.4) for *σ* = 0.3; (0.46, 1.67) for *σ* = 0.5; (0.32, 1.92) for *σ* = 0.7 (see Fig. S2).

For every lineage (population) *l* in the network, we assigned a rate *r*_*l*_ drawn from the log-normal distribution with a fixed *σ*. This rate *r*_*l*_ was applied to this lineage across all gene trees in the replicate data set, as follows.

To convert number of generations to number of substitutions per site in a gene tree edge *e*, we multiplied its length *t*_*e*_ in generations (from the coalescent simulation) by the average mutation rate *µ*, either *µ* = 1.25 × 10^*−*7^ or *µ* = 1.25 × 10^*−*8^ mutations per site per generation. In our simulations we refer to *µ* as the mutation rate, as opposed to substitution rate, because we simulated mutations in haploid genomes, that may not become fixed in a population. The slower rate 1.25 × 10^*−*8^ is similar to that in the human germline [Scally and Durbin, 2012]. The faster rate 1.25 × 10^*−*7^ is extreme for eukaryotes [Wang and Obbard, 2023, Bergeron et al., 2023] but within range for some organisms [Peck and Lauring, 2018, e.g. DNA viruses,] or relevant for studies at deeper time scales.

We then multiplied each branch length by the relative mutation rate *r*_*l*_ of the population lineage *l* that the branch evolved in, simulated as described above. Therefore the final edge length *ℓ*(*e*) of a gene tree branch *e* evolving in population *l*, in substitutions per site, was

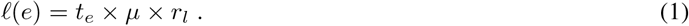

On networks other than `g1` and `g4` we used a molecular clock, that is *σ* = 0 and *r*_*l*_ = 1 always.

#### 2.1.5 Simulating sequences

Once we had 11,800 or 118,000 gene trees with lineage-specific branch lengths scaled relative to their amount of mutation rate variation, we then simulated sequences from these gene trees using `Seq-Gen` v.1.3.4 [Rambaut and Grass, 1997]. We used the HKY model with equilibrium base frequencies of 0.2 A, 0.3 C, 0.3 G and 0.2 T and a transition/transversion ratio of 3. Additionally, we simulated rate variation across sites using a discretized gamma distribution with shape *α* = 0.35 and 10 rate categories for the discretization. An *α* of 0.35 is representative of estimates from whole genomes in mammals [Arbiza et al., 2011].

To simulate a data set of unlinked sites, we aimed to extract a single variable site per gene tree. For each gene tree, we simulated an alignment of 500 base pairs. We took the first biallelic or multiallelic site in each alignment and wrote them to a VCF file. If the alignment yielded no biallelic or multiallelic sites, we re-simulated the alignment up to 20 times. We simulated alignments until we reached *n* = 10,000 or *n* = 100,000 biallelic sites.

Once we had a `VCF` file with *n* (10,000 or 100,000) biallelic sites, we generated a file mapping individual names to population names in. `ind EIGENSTRAT` format. From there, we converted the `VCF` file to `EIGENSTRAT` format using the `vcf2eigenstrat.py` Python script from Iain Mathieson, available at https://github.com/mathii/gdc/blob/master/vcf2eigenstrat.py, using our individual mapping file to specify populations. These `EIGENSTRAT` files served as input for `admixtools` analyses, detailed below.

### 2.2 Analysis of the simulated data

First, we calculated block-specific *f*_2_ values for each replicate with the `admixtoolsfunction f2_from_geno` using the `EIGENSTRAT` files as input, using default settings except for specifying blocks of size 1 site. One such default setting was `adjust_pseudohaploid = TRUE`, which ensures that samples are treated as haploid. These *f*_2_ values were used for graph inference and *f*_4_ tests below.

#### 2.2.1 Tree-tests from *f*_4_ statistics

For simulations under `g1` and `g4`, using the `f4` function with default settings and `f4mode = TRUE`, we calculated *f*_4_ statistics, their standard errors and performed the *f*_4_ tests (of *f*_4_ = 0 expected under no reticulation). For each set of 4 populations, 3 *f*_4_-tests can be performed, one for each of the 3 unrooted topologies on 4 taxa. We performed the test corresponding to the split *p*_1_*p*_2_ |*p*_3_*p*_4_ where {*p*_1_, *p*_2_} were the 2 populations that received most support by the data as being sisters. This was determined as the pair having the smallest *f*_2_ statistic, (which measures genetic divergence) among the 6 pairs from the selected 4 populations.

#### 2.2.2 Admixture graph inference

From all simulated data, we estimated admixture graphs using `find_graphs` from `admixtools`, with chimpanzee as outgroup. Search settings were chosen to match or exceed those in Maier et al. [2023]’s simulations, so as to be sure we explored graph space thoroughly, yet in a feasible timeframe (see Supplementary Material, Table S1). Specifically, settings increased from default values were as follows: number of generated and evaluated graphs per generation (`numgraphs`) to 25, total number of generations after which to stop generating graphs (`stop_gen`) to 10,000, number of generations without improvement after which to stop generating graphs (`stop_gen2`) to 60, number of generations without best score improvement before a new graph search strategy is attempted (`plusminus_generations`) to 100, and number of random initializations of starting weights (`numgraphs`) to 100.

We ran `find_graphs` for each replicate with various maximum allowed number reticulations, called “search *h*” (changing `max_admix`). For data simulated under networks with 1 reticulation, the search *h* was set to 0 (to estimate a tree), 1 (the true *h*) or 2. For data simulated under networks with 2, 3 or 4 reticulations, the search *h* was set to 0, 1 or the true *h*. For each replicate data set and each search *h*, we ran 50 independent runs of `find_graphs`, each started with its own seed. In all, `find_graph` was run 150 times for each replicate data set (3 search *h* values ×50 number of independent runs).

For each data set and each search *h*, we combined graphs across all 50 independent runs. We then filtered this large ensemble of graphs to keep a set of “top graphs”, as follows. We identified the best graph with the highest log-likelihood score, *Ŝ*, and ranked all graphs by their scores. The first 5 graphs (with the 5 best scores) were kept in the set of top graphs, as well as any graph with a score *S > Ŝ −* 10.

Keeping multiple top graphs and looking for stable topological features (see below) is in line with best practices recommended by Maier et al. [2023]. Also note that a drop threshold of 10 in log-likelihood is conservative, leading to a very large set of top graphs. For example, a threshold drop of 3 in log-likelihood, less conservative, could be argued for. It corresponds to an increase of 2 × 3 = 6 in the Bayesian information criterion (BIC) for graphs of the same *h* and same number of free edge parameters – a change traditionally considered as strong evidence to reject the graph with the worse score [Raftery, 1995].

For each replicate data set, we also calculated the log-likelihood score of the true graph topology using `qpgraph`.

### 2.3 Measures of accuracy

#### 2.3.1 Recovery of the true graph in the top set

We first calculated the network inference accuracy as the probability that the true network is part of the set of top graphs. Equality (isomorphism) between a top graph and the true network was evaluated based on equal “hash” output by `admixtools.` Given our conservative set of top graphs, this accuracy measure is quite “generous”: it does *not* require that the true network has the best log-likelihood. Rather, it requires that the true network has one of the best 5 scores, or has a score within 10 units of the best score found.

As the true `g4` topology was almost never in the set of top graphs, we looked for network features that might be correctly inferred in top graphs.

#### 2.3.2 Recovery of the true hardwired clusters

First, we checked if hardwired clusters were recovered. Each edge in a network is associated with a hardwired cluster: the set of taxa (populations) that contain genetic material inherited from that edge. For example, `g1` contains the cluster {Denisovan, Neanderthal east, Neanderthal west} from its hybrid edge with *γ* = 0.52. The *hardwired-distance d*(*N*_1_, *N*_2_) between networks *N*_1_ and *N*_2_ is the number of clusters found in one but not the other network [Huson et al., 2010]. This is the Robinson-Foulds distance when *N*_1_ and *N*_2_ are trees. More generally, *d* is not a proper distance, because distinct networks may share the same clusters. Fig. S3 shows networks *N* ≠`g4` with an identical list of clusters, such that *d*(*N*,`g4`) = 0.

#### 2.3.3 Recovery of the major (or near-major) tree

Second, we considered trees displayed by the inferred and true networks. The trees *displayed* in a network *N* are all the trees that can be obtained by keeping exactly one hybrid parent edge for each hybrid node, and deleting all other hybrid edges. One of these trees is the *major tree* of *N* : obtained by deleting from *N* any hybrid edge with inheritance weight *γ <* 0.5, thus keeping hybrid edges that contribute a majority *γ >* 0.5 of genetic material. Networks `g1`,`g4` and most others have the same major tree, with Denisovan sister to the Neanderthal populations. For each data set, we determined whether at least one top graph displayed the true major tree (of the true network used to generate the data).

For both `g1` and `g4`, the major tree is obtained by keeping a hybrid edge of weight *γ* = 0.52 (into Denisovan), almost tied with the alternative hybrid edge with *γ* = 0.48. So we considered the 2 displayed trees obtained by keeping either one of these 2 hybrid edges, and otherwise deleting any minor hybrid edge with *γ <* 0.48. These 2 *near-major* trees are the 2 trees displayed by `g1`, with Denisovan either sister to the Neanderthal populations, or sister to all Neanderthal and human populations. Compared to the other trees displayed by `g4`, the 2 near-major trees represent a greater proportion of the genome, so we expected that inferred graphs could display them more often than the other displayed trees. We calculated the number of true near-major trees displayed by each top graph (0, 1 or 2), then took the maximum over top graphs, for each data set.

#### 2.3.4 Search depth and true graph score

To check that our settings lead to a sufficiently thorough search of the graph space, for each data set we calculated the score difference Δ*S* = *S*(`g`) −*Ŝ*, where *S*(`g`) is the score of the true graph topology and *Ŝ* is the best score found during the search. When the search *h* is equal to the true *h*, the true topology is part of the space being searched, so a sufficiently thorough search should lead to Δ*S* ≥ 0 (as `admixtools` uses a score equal to the negative likelihood, up to a constant).

#### 2.3.5 Model selection using *f*_4_ worst residuals

To determine the impact of rate variation on model selection on the number of reticulations, we calculated the “worst residual” (WR) for each top graph, for simulations under `g1` or `g4`. This is obtained by comparing the *f*_4_ statistic observed from the data with that expected from the network (residual), calculating a z-score for each and then taking the worst residual z-score across all subsets of 4 populations. In most empirical studies, a graph with an absolute WR below 3 is interpreted as fitting the data adequately. Therefore, we performed the following model selection procedure for each replicate data set. We considered all top graphs obtained under a search *h* equal to the true *h* (1 for data generated under `g1`, 4 under `g4`). If at least one of these top graphs had |WR| ≤3 then we concluded that *h* (or fewer) reticulations was adequate to explain the data. Otherwise, if all these top graphs had |WR |> 3 then we concluded that this *h* could be rejected, and more reticulations were necessary to explain the data, resulting in a type-1 error. This threshold of 3 is widely used [e.g. Maier et al., 2023, Gutaker et al., 2020, Zhao et al., 2023, Ge et al., 2023]. For data generated under `g1`, we also calculated the WR of top graphs obtained under a search *h* of 2, to determine if *h* = 2 was accepted as adequate in case *h* = 1 was rejected.

### 2.4 Assessing topological identifiability

We used `qpgraph` to find edge parameters on the `g4` topology that best fit 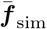, the average *f* values simulated under `g4`, averaged across the 100 replicated data set from the baseline parameters. We used option `lsqmode=2` of `qpgraph` to fix the covariance matrix *Q* to the identity, such that a network score is simply its sum of squared residuals. We extracted the fitted *f* values from this graph, ***f*** _g4_, which are theoretically expected from `g4` given the fitted edge parameters. We checked that 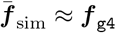, since 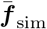 was from data simulated under `g4` — although not under the infinite-site model underlying the theory used by admixtools to calculate ***f***_g4_. We then used ***f***_g4_ as data to fit alternative topologies with qpgraph. Alternative candidate topologies consisted of all networks we used to simulate data, and the best-scoring graph from each data set simulated under `g4` and baseline parameters, if it had the same tree of blobs as `g4`.

If a topology g fits ***f*** _g4_ perfectly, then `g4` is not distinguishable from `g’` based on *f* statistics. A perfect fit corresponds to a score of 0. Due to numerical precision, we looked for graphs with a score as small as that of `g4`: *S*(`g`) *< S*(`g4`) ≈ 0. Finding one or more such topology g would show numerically that `g4` is not identifiable.

## 3 Results

### 3.1 Type-1 error and power of *f*_4_ tests

In taxon quartets with no reticulation, *f*_4_-tests correctly infer a lack of hybridization across mutation rates and number of biallelic sites in the absence of rate variation (Fig. S4, *σ* = 0 in pink) staying around the desired 5% rejection rate. Also, type-1 error stayed around 5% for 10,000 biallelic sites and a slow mutation rate (1.25 × 10^*−*8^) regardless of rate variation.

Type-1 error was moderately elevated (as high as *~* 9%) at higher levels of rate variation (*σ ≥* 0.5, blue and purple) for 10,000 biallelic sites and a fast mutation rate (1.25 *×* 10^*−*7^) or 100,000 biallelic sites and a slow mutation rate (1.25 *×* 10^*−*8^). Type-1 error was at its worst for data with 100,000 biallelic sites and a fast mutation rate, nearing about *~* 25% under our highest level of rate variation.

In scenarios with elevated type-1 error, increasing the number of individuals per population mitigated the problem. There was no clear trend of *f*_4_ performing better or worse depending on network (`g1` or `g4`).

The power of *f*_4_ to correctly detect the presence of reticulation was most affected by the data set size, unsurprisingly (Fig. S5). In all scenarios, mutation rate and rate variation across lineages had little effect. With 10,000 biallelic sites, the presence of reticulation was detected between 5% (from 1 individual/population) to 25% of the time at most (with 10 individuals/population), regardless of the number of reticulations. Larger data sets (100,000 biallelic sites) had much more power to detect reticulation, especially with more individuals per population. However, power generally decreased with more reticulations. At 1 hybridization event within a 4-population subnetwork (quarnet), power averaged 87%. At 3 hybridization events within a quarnet, average power decreased to 56%.

### 3.2 Graph inference accuracy

We obtained sets of top graphs (1 set per replicate data set and search *h* value) ranging from 5 to 16,195 distinct graphs.

As a primary measure of accuracy, we examined how many replicate data sets recovered the true network in their set of top graphs, when searching for the correct number of hybridizations.

The less complex graph `g1` was recovered for nearly all replicates, across rate variation levels and data set size when the mutation rate was low (*µ* = 1.25 ×10^*−*8^, Fig. 4 and Fig. S6). Under a faster mutation rate, rate variation across lineages started to affect accuracy. Particularly on large data sets (100,000 biallelic sites) and under a fast mutation rate (*µ* = 1.25 × 10^*−*7^), there was a large gap in accuracy between a molecular clock (100% of replicates recovered the true network in their set of top graphs) and the highest level of rate variation (49%-68% accuracy, depending on the number of individuals per population) for `g1` (Fig. 4).

**Figure 4:**
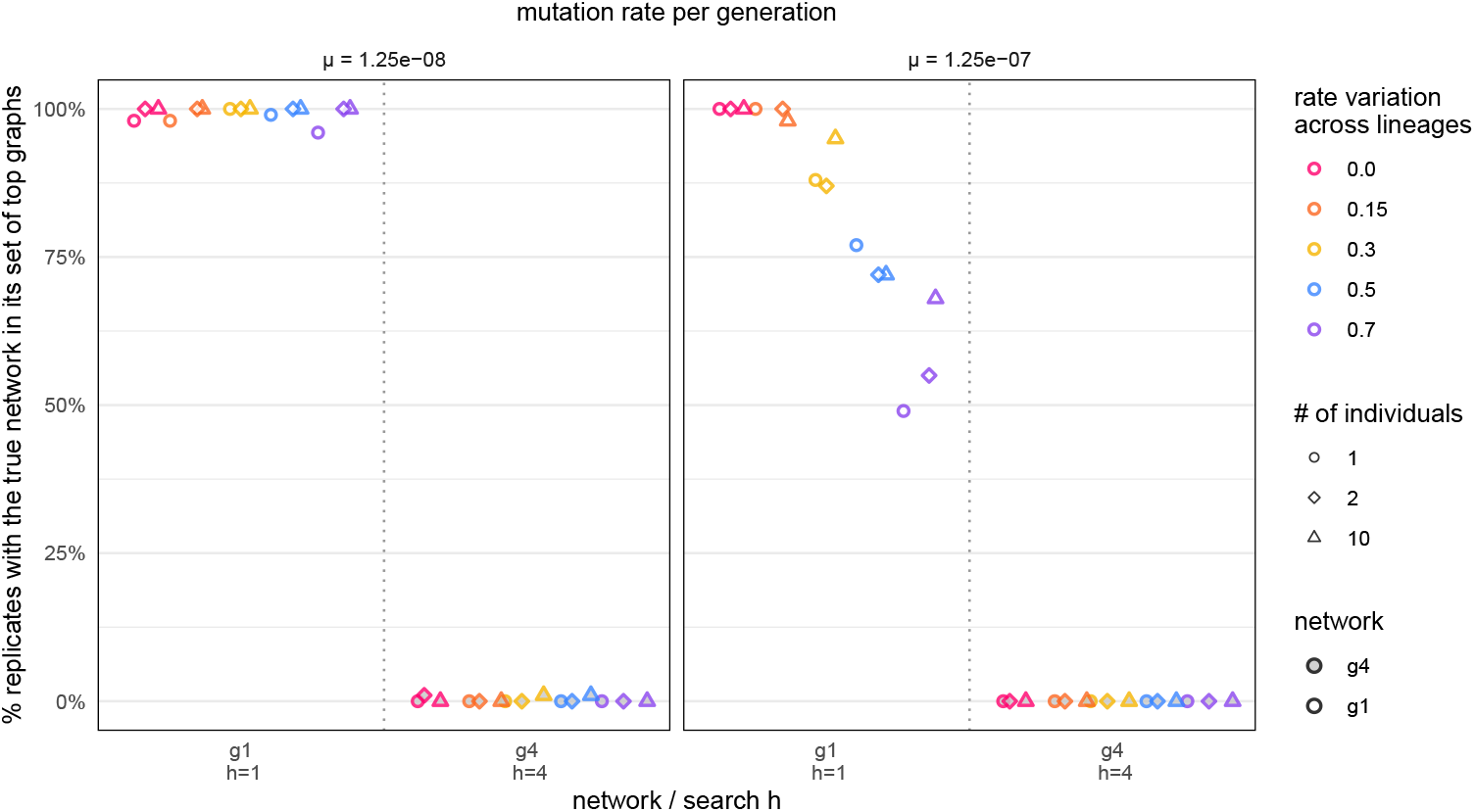
Percentage of data sets with *n* = 100,000 biallelic sites whose set of top graphs includes the true network, when the search *h* is equal to the true number of reticulations (1 under `g1`, 4 under `g4`). Panels separate different mutation rates *µ*. Color denotes the amount of rate variation (standard deviation *σ*, on a log-normal distribution with mean 1 as in Fig. S2), with pink representing *σ* = 0 (a molecular clock), orange *σ* = 0.15, yellow *σ* = 0.3, blue *σ* = 0.5, and purple *σ* = 0.7. Shape represents number of individuals per population: 1 (circles), 2 (diamonds) or 10 (triangles). Empty shapes are for data simulated under `g1` and filled shapes under `g4`. For accuracy from data with *n* = 10,000 biallelic sites, see Fig. S6.

However, `g4` was almost never recovered in the inferred set of top graphs, regardless of other factors (Figs. 4 and Fig. S6): accuracy was 0% in 56 of the 60 parameter combinations, and 1% otherwise.

Modifying the parental inheritance values (*γ*) did not make a material difference in recovering `g1` or `g4`, under baseline parameters. Accuracy remained at 100% for `g1` when the minor *γ* was decreased from 0.48 to 0.06. Accuracy increased from 0% to 2% for `g4` when its 3 smaller *γ*s were increased from 0.06, 0.19, 0.32 to 0.32, 0.48 and 0.48 respectively.

The number of hybridizations with a minimum cycle size of *c* = 3 correlated with the probability of recovering the true network among the set of top graphs (Fig. 5). Networks without such hybridizations were recovered with high accuracy, including a level-3 network recovered from 81% of replicate data sets (`g3-l3-c44`, *h* = 3). In contrast, a level-1 network with *h* = 2 reticulations both with *c* = 3 was recovered with only 6% accuracy (`g2-l1-c33`).

**Figure 5:**
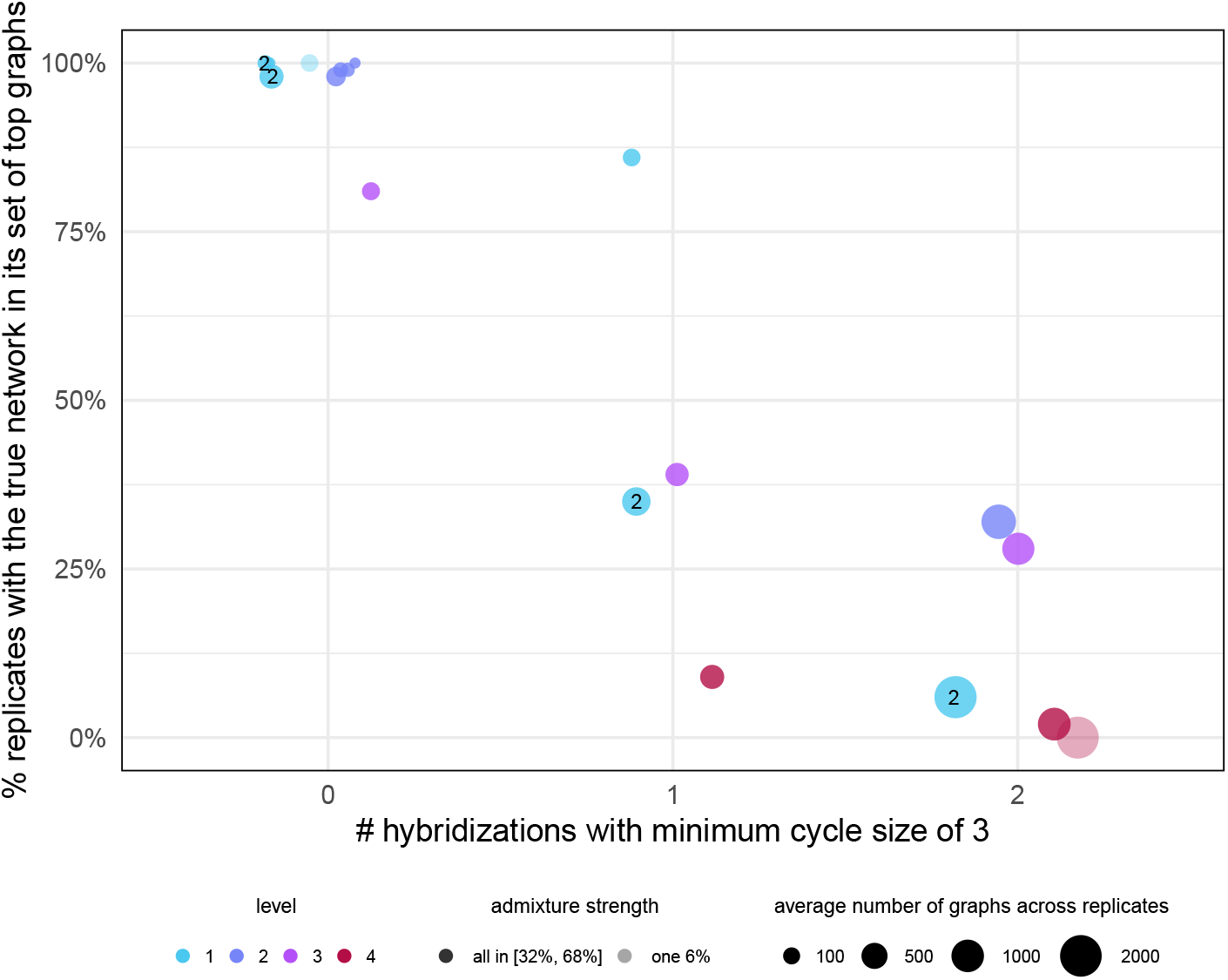
Percentage of data sets whose set of top graphs include the true network when searching for the true number *h* of reticulations, under baseline parameters: 10 individuals per population, 100,000 biallelic sites, *µ* = 1.25 × 10^*−*8^ and a molecular clock. Each point corresponds to `g1`, `g4`, or one population network in Fig. S1. Points are labeled with the number of reticulations *h* if it does not match the level *ℓ* (indicated with colors). Two networks (light points) had weak admixture weights, including one *γ* = 0.06. Point area is proportional to the average number of top graphs.

We then measured accuracy by requiring at least one top graph at hardwired-distance 0 from the true network. For a network `g` of level 1, this is equivalent to requiring that one top graph is equal to `g`. But for other networks, this is a relaxed measure of accuracy. (Fig. S3 shows an example graph that differs from `g4` but is at hardwired-distance 0 from it). Despite this relaxed measure, accuracy remained largely influenced by the number of reticulations with a minimum cycle size of 3 (Fig S7). With no such reticulations, accuracy increased to 98% or higher. On networks with 2 such reticulations, the best improvement was from 28% to 42% (for the tree-child network `g3-t334`). For `g4`, accuracy was 5% at best (Fig. S8).

Given that almost none of the top graphs were at hardwired-distance 0 from `g4`, we looked at the distance of the closest top graph to the true network, as a more informative measure of accuracy (with lower distances corresponding to better accuracy). Unsurprisingly, accuracy was best when searching for the correct *h* (Fig. 6 for `g4`, Fig. S9 for `g1`). Importantly, there was strong evidence that rate variation across lineages decreases accuracy (likelihood ratio test *p <* 10^*−*15^; see Supplemental section 5), particularly at levels *σ* ≥ 0.3. In other words, more rate variation increased the distance of the closest top graph to the true network.

**Figure 6:**
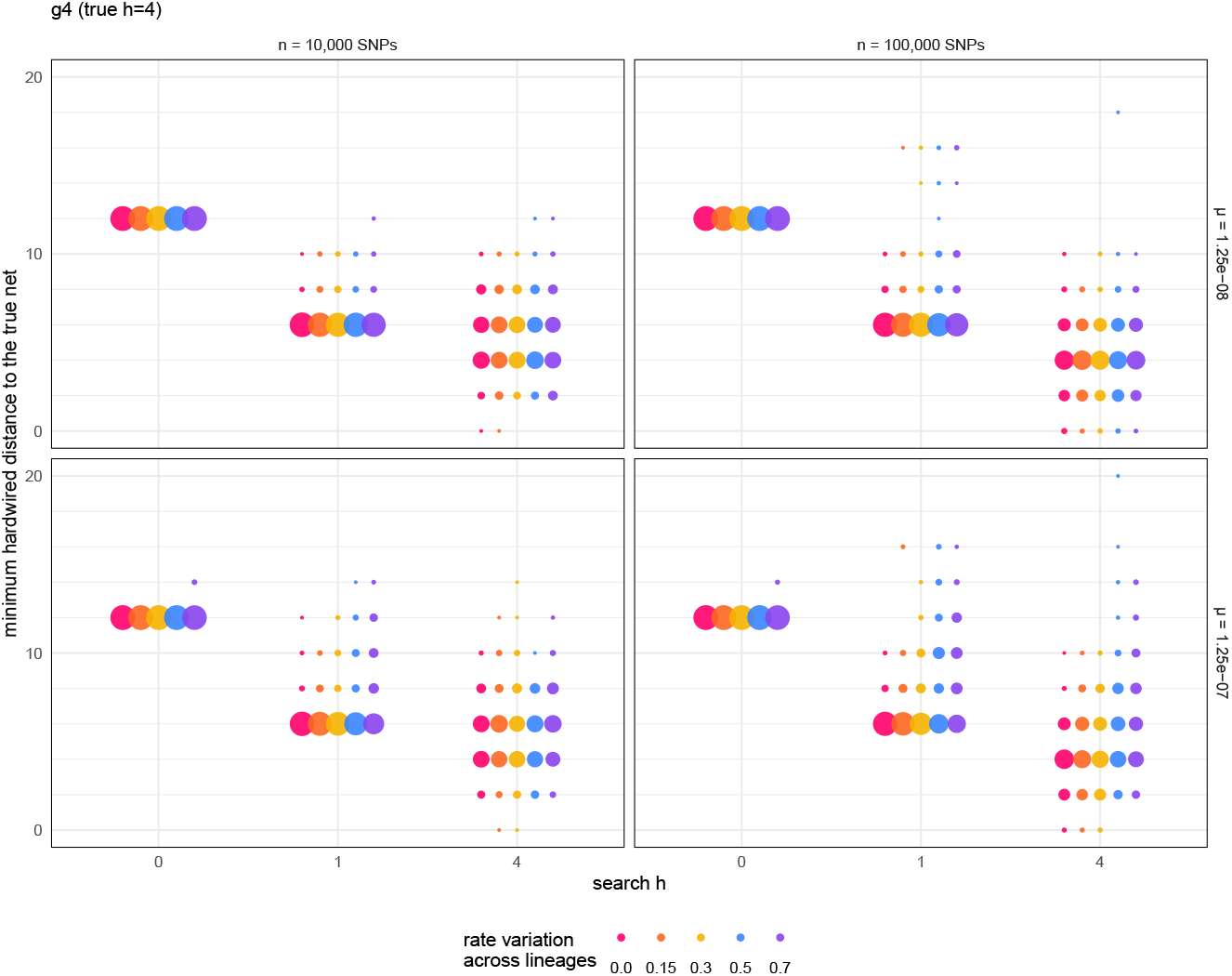
Distribution of the hardwired-distance between the true network and the closest top graph (that is, minimum hardwired-distance to the true network across top graphs), for data sets simulated under `g4`. The area of each circle is proportional to the percentage of replicate data sets, among the 300 data sets with a given combination of mutation rate *µ* and number of biallelic sites *n*. Color denotes the amount of rate variation as in Fig. 4.

### 3.3 Accuracy of graph features

As `g4` and some other networks were rarely correctly estimated, we assessed whether inferred top graphs contained graph features of the true network. We focused on two features: whether top graphs display the true network’s major tree, and whether top graphs display any (or both) of the 2 near-major trees.

#### 3.3.1 Display of the major tree

Under the best-case baseline parameters, the major tree was recovered (displayed in at least one top graph) 99.9% of the time when searching for the correct number of reticulations. Even when considering a small set of top graphs, the true major tree was displayed in the single best top-scoring graph 46% of the time, and in one of the best 5 top graphs 70% of the time overall. This high accuracy measure is in stark contrast with the low accuracy to infer the full network when 1 or more reticulations has *c* = 3.

Under `g1` and non-baseline parameters, the major tree was recovered very frequently (Fig. S10), even when searching for an underfitting *h* = 0 reticulations. When searching for trees only (*h* = 0), this metric essentially states whether the inferred tree was the major tree. When doing so, increasing the number of individuals per population had a major effect, increasing the ability to infer the major tree from as low as 25% (1 individual) to nearly 100% (10 individuals/population). Rate variation across lineages had a substantial effect on the larger data set (100,000 biallelic sites) with a fast mutation rate (*µ* = 1.25 *×* 10^*−*7^), with increased lineage-rate variation decreasing the ability to display the major tree, especially at search *h ≥* 1.

Under `g4`, the major tree was almost always recovered when searching for the correct number of 4 reticulations (Fig. S11). Even when searching for one reticulation, the major tree was recovered fairly frequently from 10,000 biallelic sites (51%-95%), but less frequently from 100,000 biallelic sites (0%-40%). Surprisingly, more individuals lead to a lower recovery of `g4`’s major tree when searching for graphs with at most 1 reticulations (Fig. S11). This may be, in part, because data with more individuals tended to have fewer top graphs than data with few individuals (see Table S2), and keeping more top graphs increases the chance that one of them displays the major tree. This reduction in the number of top graphs can be the result of our filtering criterion (keep graphs of score *S > Ŝ* −10) and the tendency of larger data sets to have larger score differences between candidate networks. When searching for trees (*h* = 0), the true major tree was recovered 0%-54% of replicates, with more individuals per population greatly improving performance (in this case, 5 top graphs were kept almost always).

Unlike for `g1`, the effect of rate variation was limited compared to the effect of other factors.

When the major tree is not recovered, we can look at the best hardwired-distance between the major tree and trees displayed by top graphs (Figs. S12 and S13). This distance is almost always at its minimum (2), showing that when the major tree is not recovered, a very similar tree is displayed by top graphs.

#### 3.3.2 Display of the two near-major trees

On `g1` and `g4`, the accuracy of displaying both near-major trees showed the same pattern as that of displaying the major tree (Figs. S14 and S15). One exception was when constraining top graphs to be trees (search *h* = 0). In that case, a top graph may display (be equal to) the true major tree, but may never display both near-major trees, clearly. In short, both near-major trees were recovered with high accuracy under `g1` and with good-to-high accuracy under `g4` (depending on the search *h*); and rate variation across lineages decreased accuracy in several conditions.

With a search *h* equal to or above the true *h*, if one top graph displayed the major tree, then some top graph was very likely to actually display both near-major trees (Figs. S16 and S17). With a search *h* = 1 and under `g4`, many data sets have their top graphs display at most 1 near-major tree, not both (Fig. S17).

### 3.4 Accuracy to select the number of reticulations

Based on the worst residual criterion to select the number of reticulations *h*, rate variation across lineages and number of biallelic sites both affected the ability to correctly choose *h* = 1 for `g1`. With smaller data sets (10,000 biallelic sites) and low levels of rate variation, “*h* ≤ 1” was rejected in about 3% of data sets (Fig. 7, *σ* ≤ 0.3 in pink, orange and yellow). For the smaller data sets, this type-1 error rate increased beyond the traditionally tolerated 5% when rate varied more across lineages and the overall mutation rate was fast (1.25 *×* 10^*−*7^)

**Figure 7:**
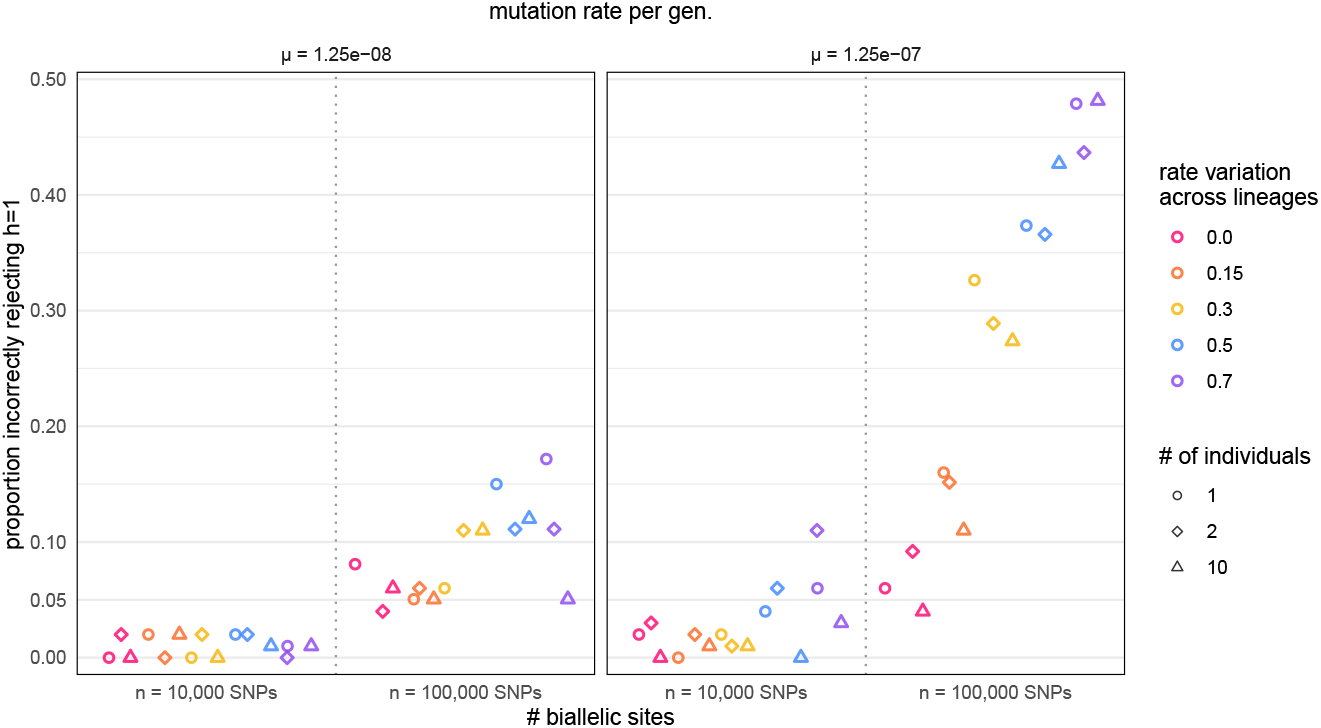
Type-1 error rate from worst residuals: proportion of data sets simulated under `g1` incorrectly rejecting *h* ≤ 1. For each data set, top graphs were obtained by searching for high-likelihood graphs with at most 1 reticulation. If all these top graphs had a worst residual |WR| *>* 3 then *h ≤* 1 was rejected. Shape and color is as in Fig. S4.

In the absence of rate variation across lineages, the rate at which *h* ≤ 1 was rejected was greater for bigger data sets (100,000 biallelic sites) than smaller data sets, between 5-10%. This type-1 error rate was strongly affected by rate variation across lineage for these bigger data sets, especially when the average mutation rate was fast. At its worst, *h* = 1 was rejected for almost ~ 50% data sets. In other words, under strong rate variation across lineages and a fast mutation rate, empirical practitioners with large data sets would have a 50% chance of wrongly rejecting the true *h* = 1 in favor of *h ≥* 2, when using |WR| *≤* 3 as a criterion of adequate fit. In all cases, *h ≥* 2 could be considered adequate, as there was always at least *h* = 2 graph with |WR| *≤* 3 per replicate.

Under `g4` and when allowing for up to 4 reticulations, one top graph had |WR|≤ 3 always, such that “*h* ≤ 4” was correctly inferred to fit the data adequately. While this is excellent model selection accuracy, it is paired with very large sets of top graphs that almost never contain the true network, raising doubts about model overfitting within the class of 4-reticulation networks.

### 3.5 Score difference

The difference between the scores of the true network and best-scoring graph was always Δ*S* ≥ 0 when searching for graphs with up to the true number of reticulations or more, suggesting a sufficiently thorough search of the graph space (Figs. S18 and S19).

Rate variation across lineages significantly increased this score gap Δ*S*. This may negatively impact model selection for methods based on likelihood scores such as AIC or BIC, as best-scoring graphs may be wrongly interpreted as fitting the data significantly better than the true network. Future work could investigate this further.

## 4 Discussion

One striking result from our study is the gap in accuracy between networks. Networks with *h* ≤ 2 and separate cycles of at least 4 nodes each were estimated with over 98% accuracy from *f*_4_ data. Among them, `g1` was estimated highly accurately even in the presence of rate variation across lineages despite the sensitivity of individual *f*_4_ tests to detect admixture. In contrast, `g4` and other complex networks were never or rarely recovered. We propose that this gap is due, at least in part, to differences in theoretical identifiability. When multiple networks are not distinguishable from one another on the basis of *f*_4_ data, they form a “terrace” of networks with the same score for a given data set, and cause an increase in the number of similarly high-scoring top graphs — as observed in our study. We found 6 different topologies indistinguishable from `g4` or from each other based on *f*-statistics (Fig. S25). The expected *f* values expected from these networks are identical to those expected from `g4` (and to the average *f* values simulated in our data under baseline conditions) for some combination of edge drift and inheritance parameters.

### 4.1 Identifiability of complex graphs from *f*-statistics

We are unaware of any theoretical study of topology identifiability from *f*-statistics. Maier et al. [2023] proposed a numerical tool to determine the number of free edge parameters on a given network topology, and stated that the network root is not identifiable. Consequently, inference from *f* statistics should aim to estimate a *semidirected network*, rather than a rooted network [Solís-Lemus and Ané, 2016, Xu and Ané, 2023]. Yet, there is a lack of graph theory to characterize outgroup constraints on the rooted network that do not overconstrain the semidirected network.

However, there is growing knowledge about network identifiability from other data types [Solís-Lemus and Ané, 2016, Baños, 2019, Gross et al., 2021, Allman et al., 2022, 2023, Xu and Ané, 2023, Allman et al., 2024, Rhodes et al., 2025, Englander et al., 2025]. Most similar to *f*-statistics are average genetic distances, because *f*_2_ statistics match distances on trees (and *f*_3_, *f*_4_ values are linear combinations of *f*_2_ statistics). As from *f*-statistics, the lengths of the 3 edges incident to a hybrid node are not identifiable from each other, based on average distances and under a model without incomplete lineage sorting [Xu and Ané, 2023]. This local lack of edge length identifiability is the reason to constrain hybrid edge lengths to 0 in `qpgraph.` Xu and Ané [2023] proved that many local topological features (subgraphs) are not distinguishable from one another from average distances. For example, a reticulation in a (undirected) cycle of 4 edges can perfectly reproduce the expected distance data that are expected from various subgraphs in which the cycle is modified with additional reticulations. If such features also lack identifiability from *f*_4_ statistics, then one would expect an exponential number of graphs to fit a given data set, all with a similar likelihood score, when searching for top-scoring graphs with an increasing allowed number of reticulations. This is indeed what we observed in our study.

On the positive side, studies about network identifiability found that the network’s *tree of blobs* is identifiable from many data types [Allman et al., 2023, Xu and Ané, 2023, Rhodes et al., 2025]. The tree of blobs is obtained from the network by contracting each blob into a single node. Doing so leads to a tree, whose edges are the edges in the original network that, when removed, disconnect the graph. For `g4` for example, the tree of blobs is almost reduced to a star tree, except for a non-African clade. In contrast, `g1`’s tree of blobs retains the modern human subtree and the Neanderthal clade.

Also identifiable from many data types are networks of level 1 without “small” cycles, with “small” depending on the data type, typically with at most 3 nodes [Baños, 2019, Gross et al., 2021, Xu and Ané, 2023, Allman et al., 2024]. Given that 3-node cycles cannot be detected from some data and models, we looked at the presence of such cycles in networks. If level-1 networks are identifiable from *f* statistics as well, this may explain the accuracy of estimating level-1 networks without 3-cycles (like `g1`) in our study.

Some recent theory points at subclasses of tree-child networks as good candidates to ensure identifiability [Allman et al., 2025, Englander et al., 2025]. In Allman et al. [2025], the tree-child networks can be of any level, but must be galled and cannot have *c* ≥ 3 for any reticulation. In Englander et al. [2025], the tree-child networks must be of level at most 2 and some may have *c* = 3 for one of two reticulations, but the full network may not have any 3-cycle. Given that even level-2 networks are generally not identifiable from many data types [Xu and Ané, 2023, Rhodes et al., 2025], we suspect that identifiablity from *f*_4_ statistics does not hold either.

We are not aware of any theoretical work considering a reticulation’s minimum cycle size, beyond level-1 networks. This quantity may be helpful in future work to determine the difficulty of identifying (in theory) or estimating (in practice) reticulations in a network.

Our findings suggest that the major tree is one graph feature that might be identifiable from *f*-statistics. However, we are unaware of any study exploring the identifiability of trees displayed by the network, regardless of data type and evolutionary model. The notion of major tree requires that one of the two (or more) parents of each admixed population be designated as major and the other one(s) as minor, based on which parental population contributed most to the admixed population’s genome. Identifiability of the major tree may be facilitated by an imbalance between parental contributions, to make the major parent easier to detect.

Theoretical work on identifiability from *f*-statistics is needed, and could borrow ideas and tools developed for other data types. Empirical analyses will benefit from knowledge of which network classes guarantee identifiability and which network features are identifiable. These features should be summarized across top-scoring graphs, as they are likely to be conserved and receive high support.

### 4.2 Recommendations for best practices

Our complex graph `g4` was almost never recovered from `find_graphs:` neither as the best-scoring graph nor in the set of top graphs, despite a quite lenient threshold for retaining graphs. With our threshold of keeping graphs within 10 log-likelihood points of the best-scoring graph, we kept a median of 1,868-6,760 graphs per replicate when searching for 4 hybridization events. Thus, the recommendation from Maier et al. [2023] to review a set of best scoring graphs is necessary when using `find_graphs` to infer complex topologies, but seems insufficient with *h* as low as 4. The adequate number of top-scoring graphs seems to be low when searching for trees (*h* = 0) and to increase drastically as more reticulations are allowed, but seems to depend on the complexity of true underlying network.

Evaluating only the single best-scoring graph, dramatically lowered accuracy for ‘simple’ networks, as did retaining only the 5 best scoring graphs (Supplemental section 9), especially when a reticulation event had a small parental inheritance (*γ*). Accuracy dropped moderately when keeping graphs scoring within 3 log-likelihoods of the best graph (instead of 10): not surprisingly, downsizing the number of top scoring graphs comes at the the cost of accuracy.

Many studies published graphs as or more complex than `g4`, inferred using `admixtools` or `AdmixtureBayes` (for example, Fig. 3 in Lazaridis et al., 2014, *h* = *ℓ* = 4; Fig. 3 in Moreno-Mayar et al., 2018, *h* = *ℓ* = 6; Fig. 5 in Narasimhan et al., 2019, *h* = *ℓ* = 12). Our results should discourage the assumption that keeping a large set (e.g. thousands) of graphs ensures that the true network is included, especially if inferred graphs are of high level. In fact, the number of graphs retained when keeping those within 3 (Fig. S22) or 10 log-likelihood points of the best graph correlated negatively with accuracy. Like Maier et al. [2023], we recommend summarizing features shared by top-scoring graphs, and report features that are stable across graphs.

Individually, *f*_4_-tests were found to be error-prone, with type-1 error exacerbated by a combination of lineage-rate variation and increased mutation rate. This is most likely due to homoplasies [Frankel and Ané, 2023, Koppetsch et al., 2024]. While graph inference is more robust to rate variation, each *f*_4_-statistic is associated with one residual. Like individual *f*_4_-tests, the worst residual (worst-fitting *f*_4_) and associated selection of the appropriate number of hybridizations is strongly affected by rate variation. We recommend assessing the extent of rate variation across lineages in empirical data sets, and using caution in interpreting *f*_4_-tests and worst residuals if rate variation is found to be strong.

In their protocol, Maier et al. [2023] recommend using top graphs identified by fitting *f*_4_ data as candidate networks to be evaluated by methods that use other data types, and that might be more computationally intensive. Our study suggests using these candidate networks with caution, as the true network may still be excluded from this candidate list. Candidate networks could be used to seed the search made by other methods that explore the network space, for example.

### 4.3 Limitations and future directions

One limitation of our work is that we focused on the inference method `find_graphs`. This was due to its popularity in high-impact papers (Fig. 1), and its computational speed. However, many alternative methods can also be used to infer population history and admixture. The faster methods make use of 4-taxon data summaries like `find_graphs` does, such as quartet concordance factors [SNaQ, Solís-Lemus and Ané, 2016], 3-taxon subtrees of rooted gene trees [`PhyloNet-MPL`, Yu and Nakhleh, 2015] or site pattern frequencies across 4-taxon subsets [`PhyNEST`, Kong et al., 2025]. Methods using data jointly across all populations should be able to distinguish between more pairs of networks. Unfortunately such methods are less used or cannot search the network space due to their computational burden. For example, `legofit` and `momi` [Rogers, 2019, Kamm et al., 2020] use the multi-population site frequency spectrum from biallelic sites. Beyond patterns at individual sites, `moment-DL` uses site position information via linkage disequilibrium statistics between pairs of sites and pairs of populations [Ragsdale and Gravel, 2019]. These statistics can be seen as extending single-locus *f*-statistics, and showed power to discriminate between complex histories [Ragsdale et al., 2023a]. Finally, methods making stronger assumptions (such as a molecular clock like `BPP` Thawornwattana et al. 2023) can also distinguish more networks, such as networks with 3-cycles, at the cost of inferring extra reticulations if their model assumptions are violated [Flouri et al., 2022].

While this work and others [Frankel and Ané, 2023, Koppetsch et al., 2024] raise concerns when the molecular clock is violated, methods to quantify rate variation across lineages and across loci are underdeveloped in the presence of reticulation [but see Snir et al., 2012, Duchêne et al., 2014, 2024, in a tree context]. The “robust” ABBA-clustering test by Koppetsch et al. [2024], available in Dsuite [Malinsky et al., 2021], may prove useful to detect introgression amongst lineages where the assumption of a molecular clock may not hold true.

Selecting the appropriate number of reticulations is particularly important. For this, the worst residual is commonly used in the admixture graph literature. Given the sensitivity of this criterion to rate variation, more robust methods are needed to select the graph complexity. Yu et al. [2014] proposed *k*-fold cross-validation, implemented in `PhyloNet-ML`, and Rogers [2019] proposed a bootstrap-based model selection, available in `Legofit.` These procedures are rarely used however, likely due to their computationally burden.

Finally, there is a lack of methods to summarize a set of networks. Such methods are needed to automatically extract features shared by a large number of high-scoring graphs, as recommended by Maier et al. [2023]. Some concepts of consensus networks have been developed [Huber et al., 2023, for level-1 networks] but more methods are needed to summarize various network features, especially those known to be identifiable and recoverable with high support.

## Data availability

The code to reproduce all analyses and figures is available at https://github.com/laufran/f4-ratevar.

## Acknowledgments

We thank anonymous reviewers for their insightful feedback, that lead us to explore more aspects of network complexity.

## Supplemental material

### 1. Simulation networks and parameters

**Figure S1:**
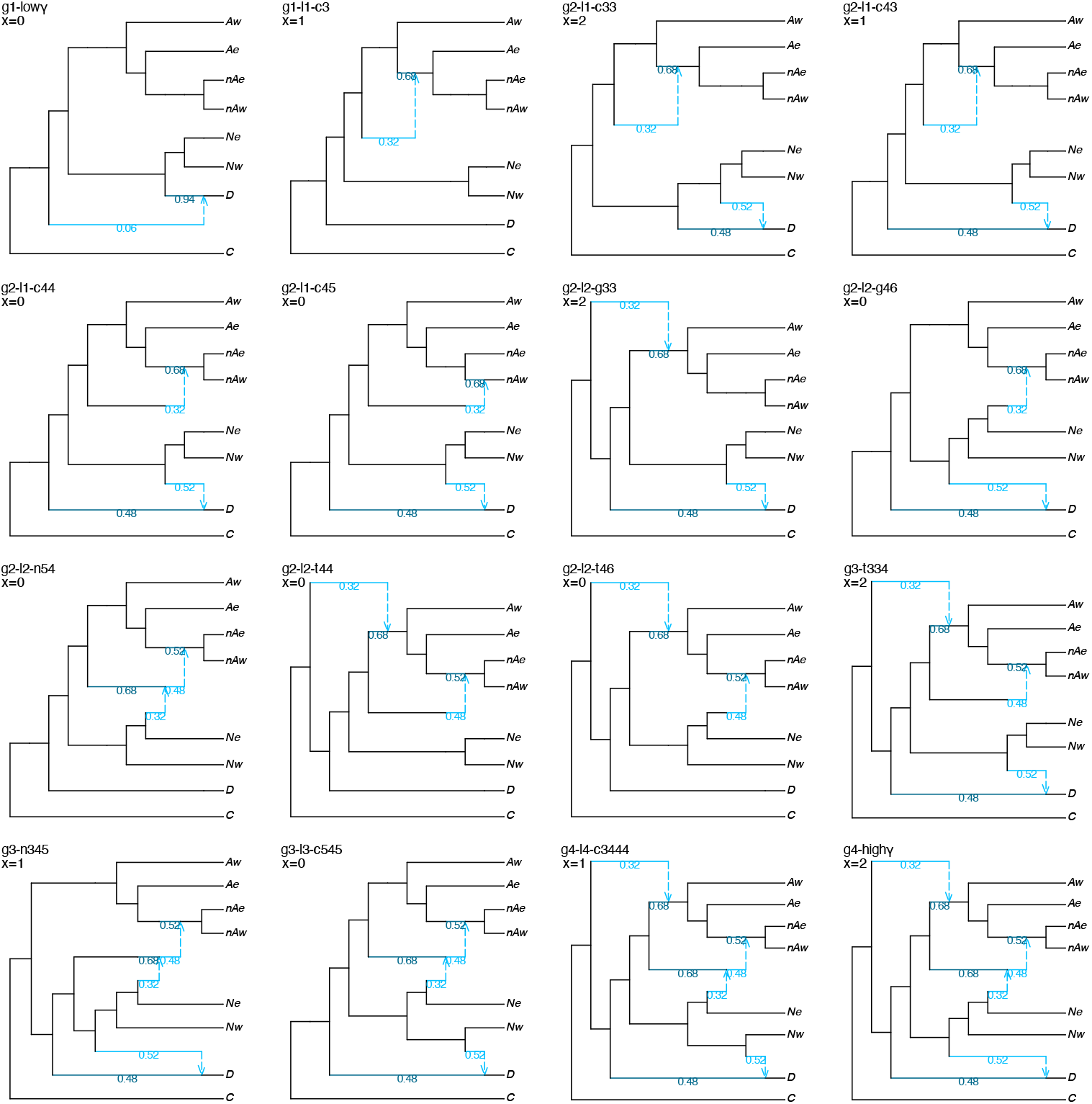
Networks used to simulate data under baseline parameters, in addition to `g1` and `g4`. In each panel, *x* denotes the number of reticulations in the network whose minimum cycle size is 3, used on the x-axis of Fig. 5. Population abbreviations: C for chimp, D for Denisovan, N for Neanderthal, A for Africa, nA for non-Africa, with suffix e or w for east and west populations. Hybrid edges are annotated with their inheritance *γ* values.

#### Population networks used to simulate data

In addition to `g1` and `g4`, networks in Fig. S1 were used to simulate data under baseline (best-case) parameters: 10 individuals per population, 100,000 biallelic sites, *µ* = 1.25 × 10^*−*8^ mutations per site per generation, and a molecular clock.

In Fig. S1, the first (top left) and last (bottom right) networks have the same topologies as `g1` and `g4` and same edge lengths, but different inheritance probabilities: low (*γ* = 0.06) in `g1-low`*γ* or high (all *γ ≥* 0.32) in `g1-high`*γ*.

For the other networks, the names indicate their number of reticulation *h* (number after `g`), their level *ℓ* (number after `l`), and some complexity features. After g*h*-l*ℓ*-, the letter `c` is for level-1 networks in which each reticulation is in a single *cycle* not affected by the presence or removal of other reticulations; `g` is for *galled* networks (but not tree-child); t is for *tree-child* networks (but not galled); and `n` is for *neither* galled nor tree-child. Numbers that follow give the minimum cycle size *c* of each reticulation in the network. `g1` is of level 1, therefore galled and tree-child. `g4` is of level-4, neither galled nor tree-child, and its hybridizations have minimum cycles sizes *c* = 3, 3, 4, 4.

Fig. 3 uses `g2-l2-g33` to illustrate the calculation of a reticulation’s minimum cycle size *c*. In section 10, `g4-l4-c3444` is shown to be non-distinguishable from `g4` based on *f*-statistics.

#### Distribution of rates across population lineages

Rate variation across lineages was simulated under `g1` and `g4.` The molecular clock corresponds to no rate variation, with *R* = 1 always.

**Figure S2:**
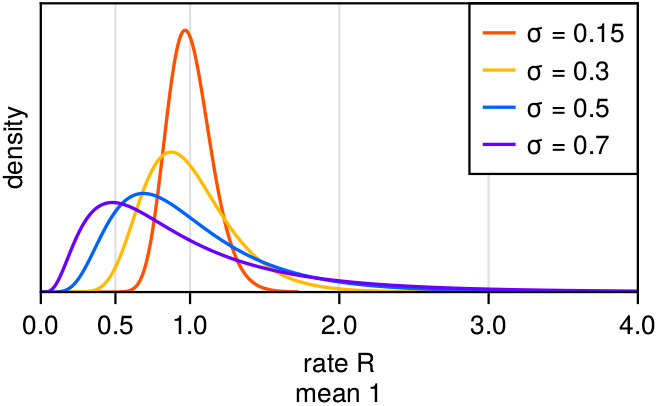
Density of the log-normal distribution used for simulating rates *R*, with standard deviation *σ* of 0.15, 0.3, 0.5 and 0.7 on the log-scale. To set the mean of *R* to 1, the mean of log(*R*) was set to *−σ*^2^*/*2.

### 2. Network topologies with identical hardwired clusters

**Figure S3:**
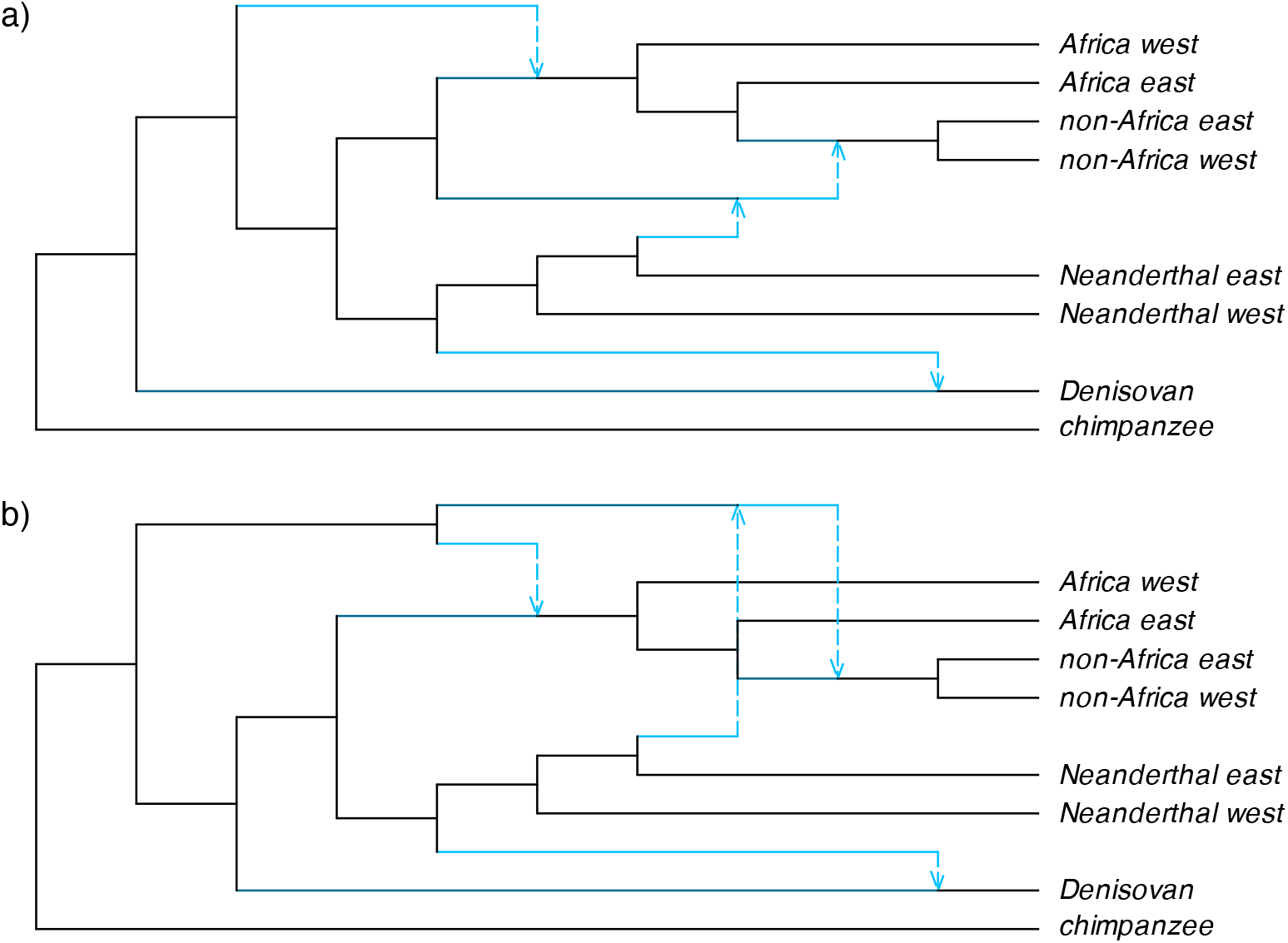
Example networks at hardwired-distance 0 from `g4`, yet different from `g4`. This example illustrates why inferred networks may be at hardwired-distance 0 of the true network g more frequently than they are identical to g (based on identical hashes). a) Network that differs from `g4` by its order of the first two divergence events: the split leading to the Denisovan lineage (on the major tree) occurring before the split with the ghost population that contributed gene flow to the 4 modern human populations (instead of after in `g4`). b) Network that differs from `g4` by the second ghost population (into non-Africans): stemming from the first ghost lineage instead of stemming from the human stem in `g4`.

### 3. Type-1 error and power of the *f*_4_-statistic to detect reticulations

**Figure S4:**
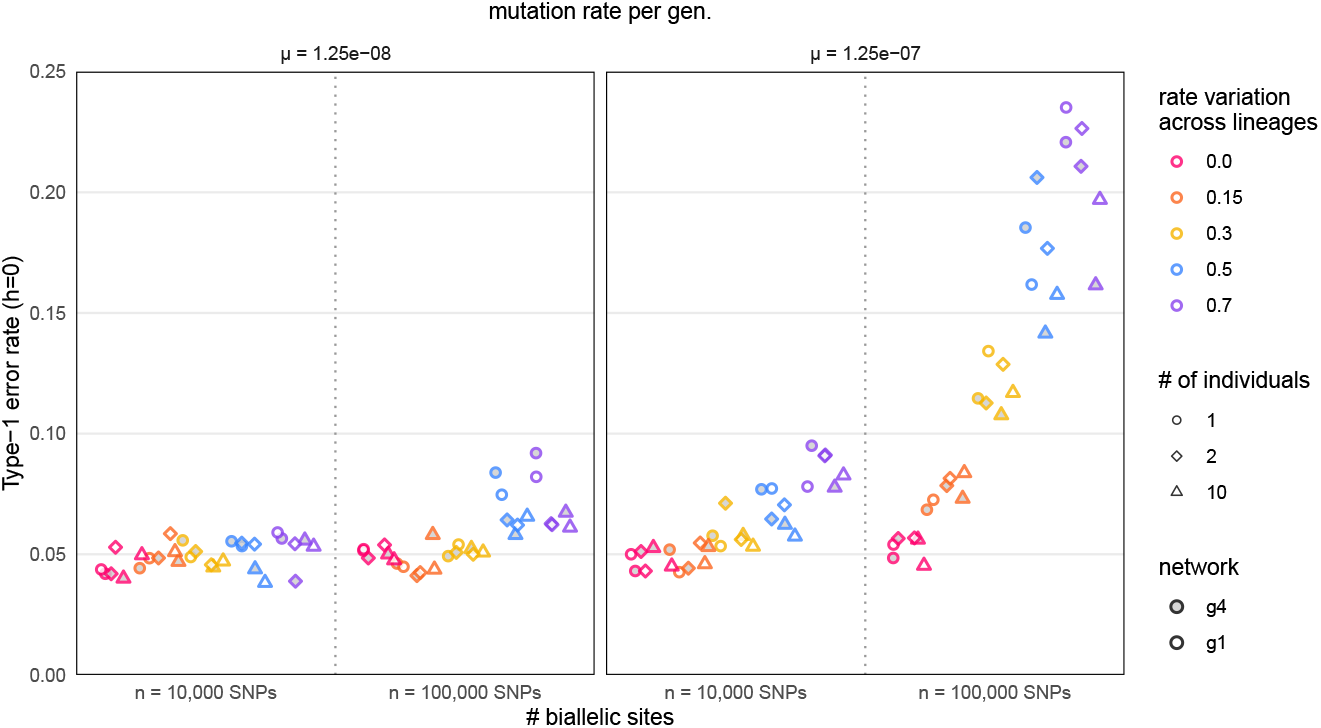
Type-1 error rate from *f*_4_: proportion of quartets incorrectly inferred to have hybridization when none was present, depending on mutation rate *µ* (1.25 × 10^*−*8^ or 1.25 ×10^*−*7^) and number of biallelic sites (10,000 or 100,000). Each point represents the proportion of tests with a p-value below 0.05 across quartets from 100 replicate data sets, with 2,600 quartets per point from the `g4` network, and 6,200 quartets per point from the `g1` network. Color denotes the amount of rate variation (standard deviation *σ*, on a log-normal distribution with mean 1 as in Fig. S2), with pink representing *σ* = 0 (a molecular clock), orange *σ* = 0.15, yellow *σ* = 0.3, blue *σ* = 0.5, and purple *σ* = 0.7. Shape represents number of individuals per population: 1 (circles), 2 (diamonds) or 10 (triangles). Shape fill indicates the network used to simulated data, either `g1` (empty shapes) or `g4` (filled shapes).

**Figure S5:**
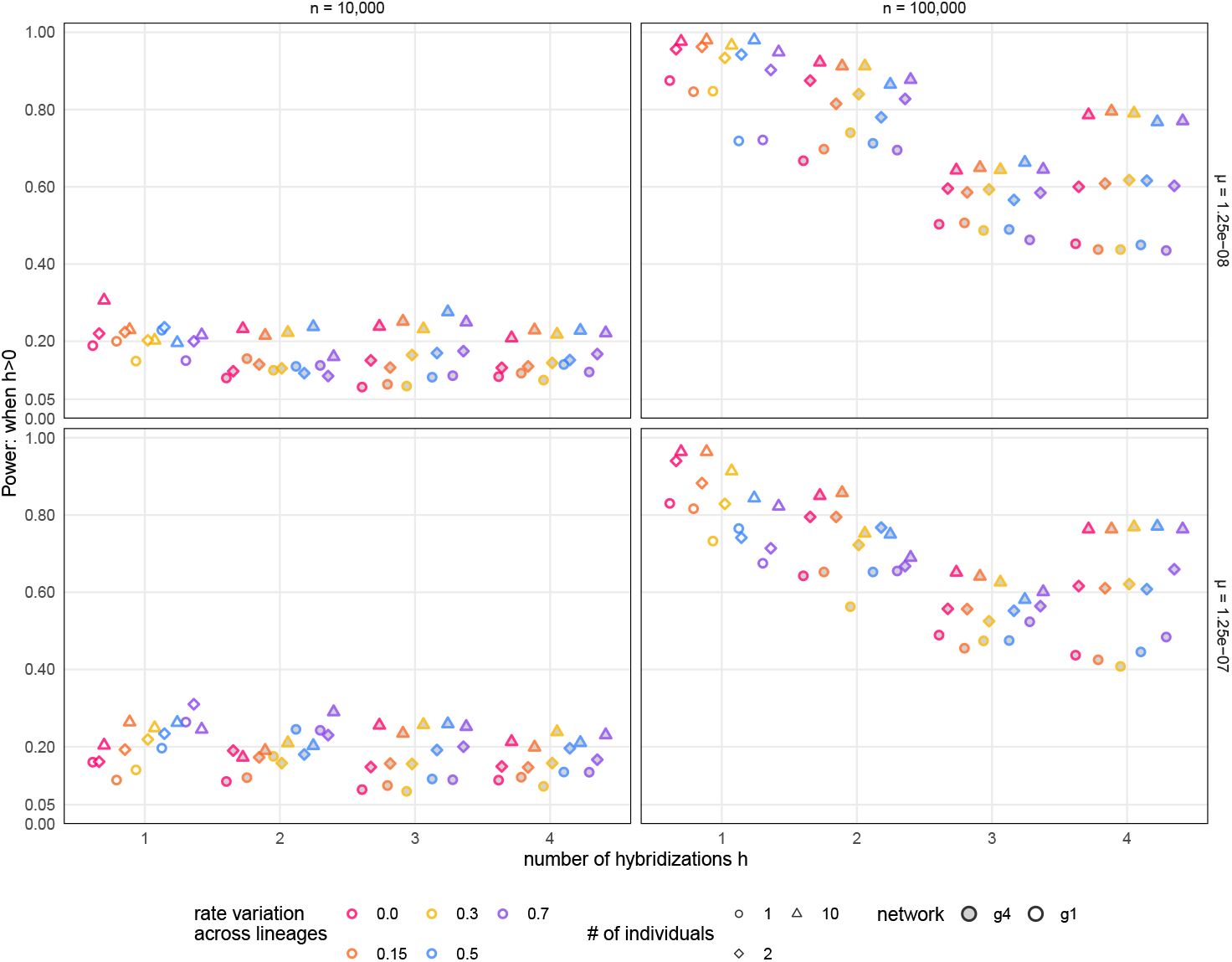
Power of the *f*_4_ statistic to detect reticulations in a given quartet. depending on mutation rate *µ* and number of biallelic sites *n*. Shape color, type, and fill are as in Fig. S4. Note that subsets of 4 populations whose subnetwork retains a reticulation have *h* = 1 from `g1` (which only has 1 reticulation in the entire network) and *h ≥* 2 from `g4`.

### 4. Complexity of graph search using find_graphs

We summarized the time needed to estimate admixture graphs using `find_graphs,` using the search parameters as specified in the Methods, for one combination of simulation parameters: across the 100 data sets simulated under `g4` (because searching for 4 admixture events is more time intensive than the maximum of 2 events with `g1`), 10 individuals per population, *n* = 100,000 biallelic sites, a molecular clock (*σ* = 0.0), and a low mutation rate of 1.25 *×* 10^*−*8^.

**Table S1:** Time needed to run find_graphs from one set of simulation parameters. Our analysis includes 50 independent runs for each of 100 data sets and each search *h*. Thus, each row summarizes 5,000 runs. The total time across all runs, search *h* values, and data sets was 474.54 days.

| search $h$ | elapsed time per run (minutes) | | | total for 1 parameter combination<br>(100 replicates × 50 runs) |
| --- | --- | --- | --- | --- |
|  | average | minimum | maximum |  |
| 0 | 2.47 | 0.779 | 5.47 | 8.57 days |
| 1 | 11.8 | 1.49 | 69.2 | 41.10 days |
| 4 | 122 | 2.67 | 697 | 424.87 days |

To ease the computational burden of running `find_graphs` across the 60 combinations of simulation parameters used for `g1` and `g4`, we ran these analyses on 13 clusters with a total of 652 threads available for parallelization. Despite this parallelization of computations, it took over 3 months (clock time) to run all these analyses. Of note, the computing time reported in Table S1 does not include the time required to calculate worst residuals, as those were calculated separately (using `qpgraph,` after determining the sets of top graphs using `find_graphs`).

For simulations on `g1` and `g4`, a total of 40,538,966 graphs were saved across all `find_graphs` runs (1 set per replicate data set and search h value). The size of sets of top graphs varied substantially across parameter settings, as shown below.

**Table S2:** Number of top graphs saved, per replicate data set and search *h* parameter, after pooling graphs from the 50 independent runs and filtering to keep at least 5 graphs, and all those with a score *S > Ŝ* −10, where *Ŝ* was the score found across the 50 runs. Each row summarizes the number of top graphs across 500 replicate data sets (100 for each level of rate variation across sites).

| search<br>$h$ | # ind<br>/pop | # biallelic<br>sites | data generated under g1 | | | | data generated under g4 | | | |
| --- | --- | --- | --- | --- | --- | --- | --- | --- | --- | --- |
|  |  |  | minimum | median | average | maximum | minimum | median | average | maximum |
| 0 | 1 | 10,000 | 5 | 5 | 5.01 | 9 | 5 | 5 | 5.43 | 15 |
| 0 | 1 | 100,000 | 5 | 5 | 5 | 5 | 5 | 5 | 5 | 5 |
| 0 | 2 | 10,000 | 5 | 5 | 5 | 5 | 5 | 5 | 5.02 | 15 |
| 0 | 2 | 100,000 | 5 | 5 | 5 | 5 | 5 | 5 | 5 | 5 |
| 0 | 10 | 10,000 | 5 | 5 | 5 | 5 | 5 | 5 | 5 | 5 |
| 0 | 10 | 100,000 | 5 | 5 | 5 | 5 | 5 | 5 | 5 | 5 |
| 1 | 1 | 10,000 | 5 | 144 | 176 | 428 | 5 | 166 | 189 | 957 |
| 1 | 1 | 100,000 | 5 | 11 | 32.5 | 133 | 5 | 8 | 12.0 | 139 |
| 1 | 2 | 10,000 | 5 | 133 | 157 | 350 | 5 | 130 | 139 | 574 |
| 1 | 2 | 100,000 | 5 | 10 | 24.4 | 129 | 5 | 6 | 7.64 | 37 |
| 1 | 10 | 10,000 | 5 | 129 | 141 | 351 | 5 | 56.5 | 89.5 | 359 |
| 1 | 10 | 100,000 | 5 | 9 | 23.3 | 132 | 5 | 5 | 5.40 | 17 |
| 2 (g1) or 4 (g4) | 1 | 10,000 | 22 | 2,112 | 1,991 | 4,433 | 183 | 6,760 | 6,755 | 16,195 |
| 2 (g1) or 4 (g4) | 1 | 100,000 | 5 | 635 | 658 | 2,637 | 29 | 2,398 | 2,363 | 6,235 |
| 2 (g1) or 4 (g4) | 2 | 100,000 | 16 | 2,120 | 1,948 | 4,517 | 245 | 5,442 | 5,565 | 11,756 |
| 2 (g1) or 4 (g4) | 2 | 100,000 | 5 | 634 | 620 | 2,636 | 5 | 2,024 | 1,955 | 5,569 |
| 2 (g1) or 4 (g4) | 10 | 10,000 | 30 | 2,158 | 2,007 | 4,679 | 728 | 5,436 | 5,459 | 12,757 |
| 2 (g1) or 4 (g4) | 10 | 100,000 | 5 | 641 | 641 | 3,012 | 28 | 1,868 | 1,854 | 4,229 |

### 5. How close are the top graphs from the true network? Recovery of the true graph from *n* = 10,000 biallelic sites

**Figure S6:**
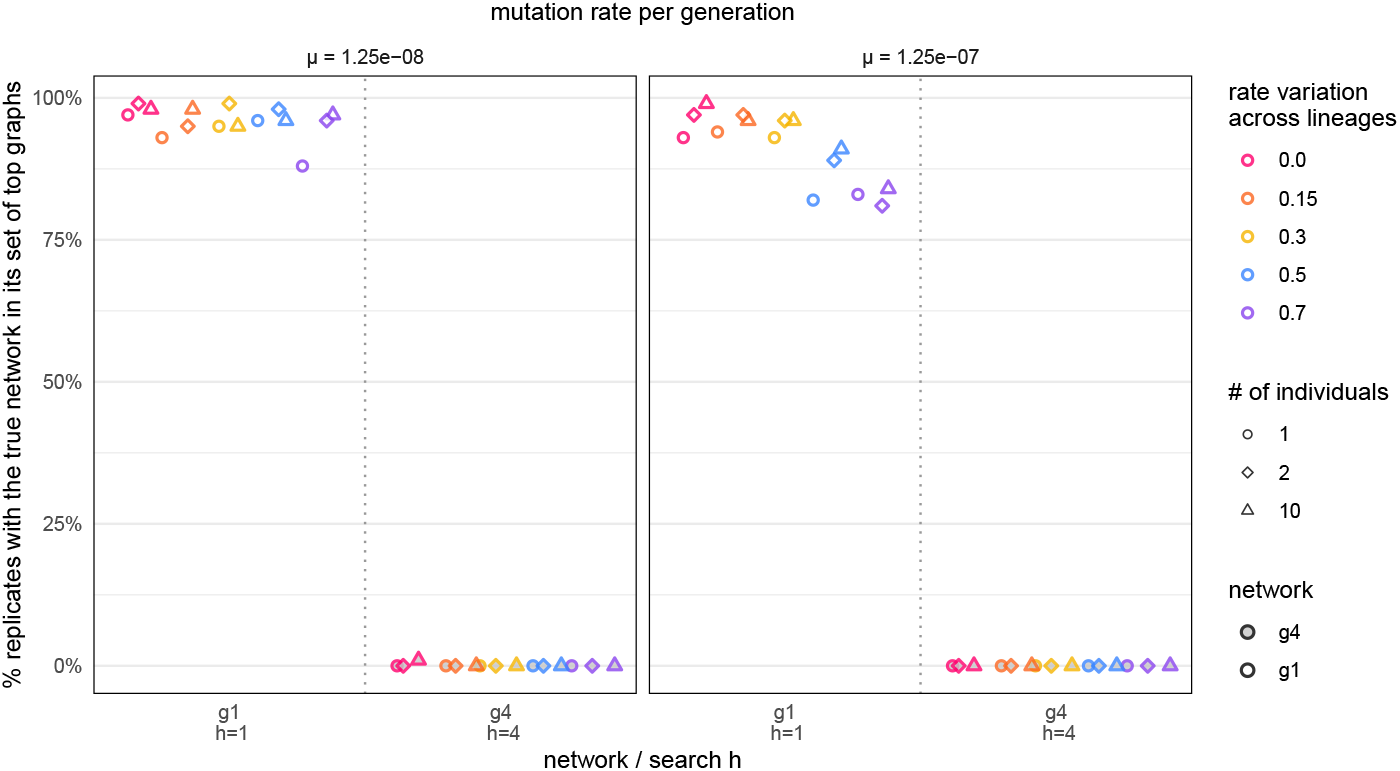
Percentage of data sets with 10,000 biallelic sites whose set of top graphs includes the true network, when searching for the true number of reticulations (*h* = 1 under `g1`, *h* = 4 under `g4`). Shape color, type, and fill are as in Fig. S4.

#### Probability that the closest top graph is at distance 0 from the true network

We calculated the hardwired-distance *d*(*G*, `g`) between each top graph *G* and the true network `g` (`g1` or `g4`), and found the closest top graph, that is, the top graph minimizing this distance: *d*_min_ = min{*d*(*G*, `g`) : *G* in top graph set}.

**Figure S7:**
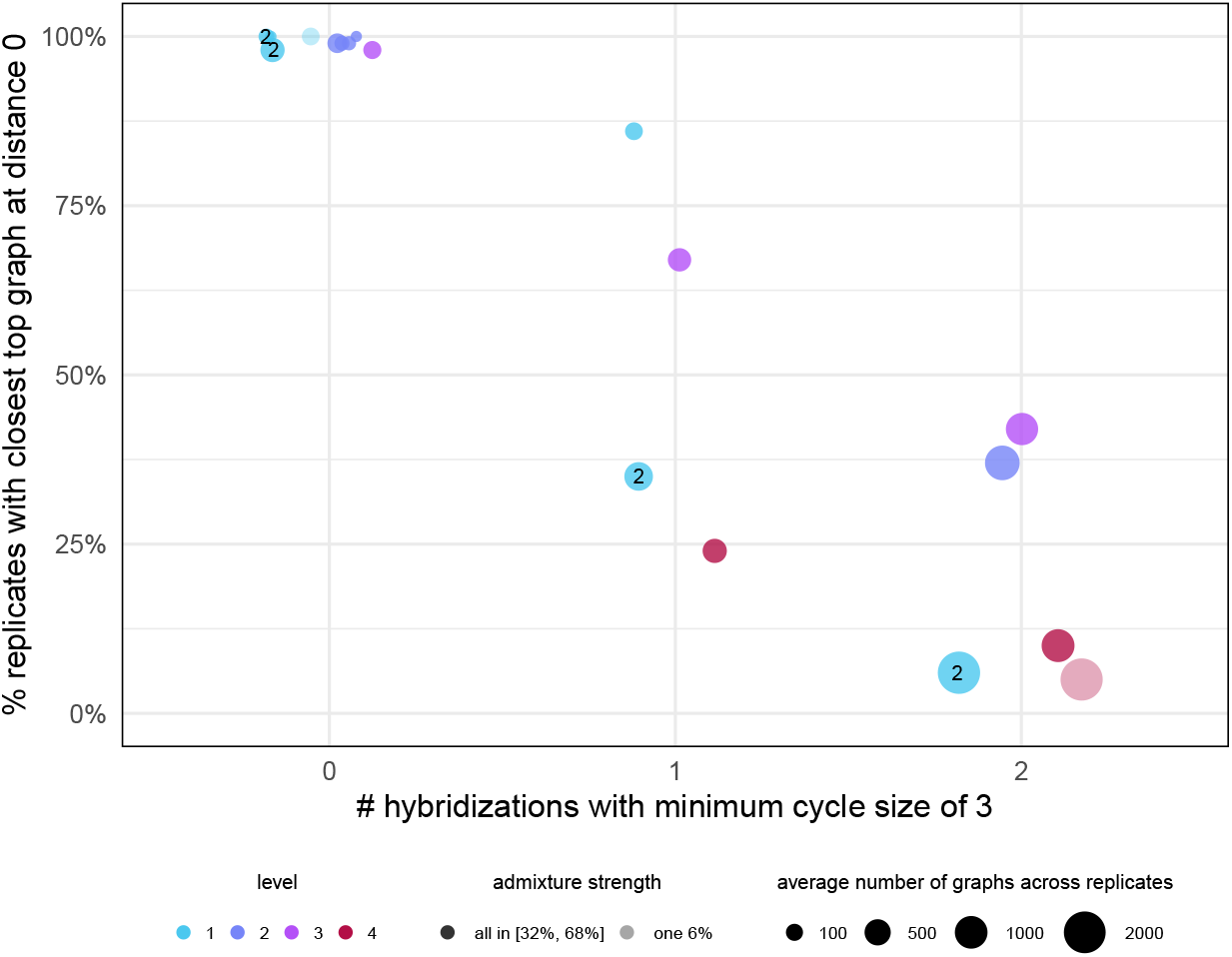
Percentage of data sets that have at least one top graph at hardwired-distance 0 from the true network when searching for the true number of reticulations *h*, under population networks in Fig. S1 and baseline parameters: 10 individuals per population, 100,000 biallelic sites, *µ* = 1.25*×* 10^*−*8^ and a molecular clock. Points are labeled with the number of reticulations *h* if *h > ℓ*. Point area is proportional to the average number of top graphs.

**Figure S8:**
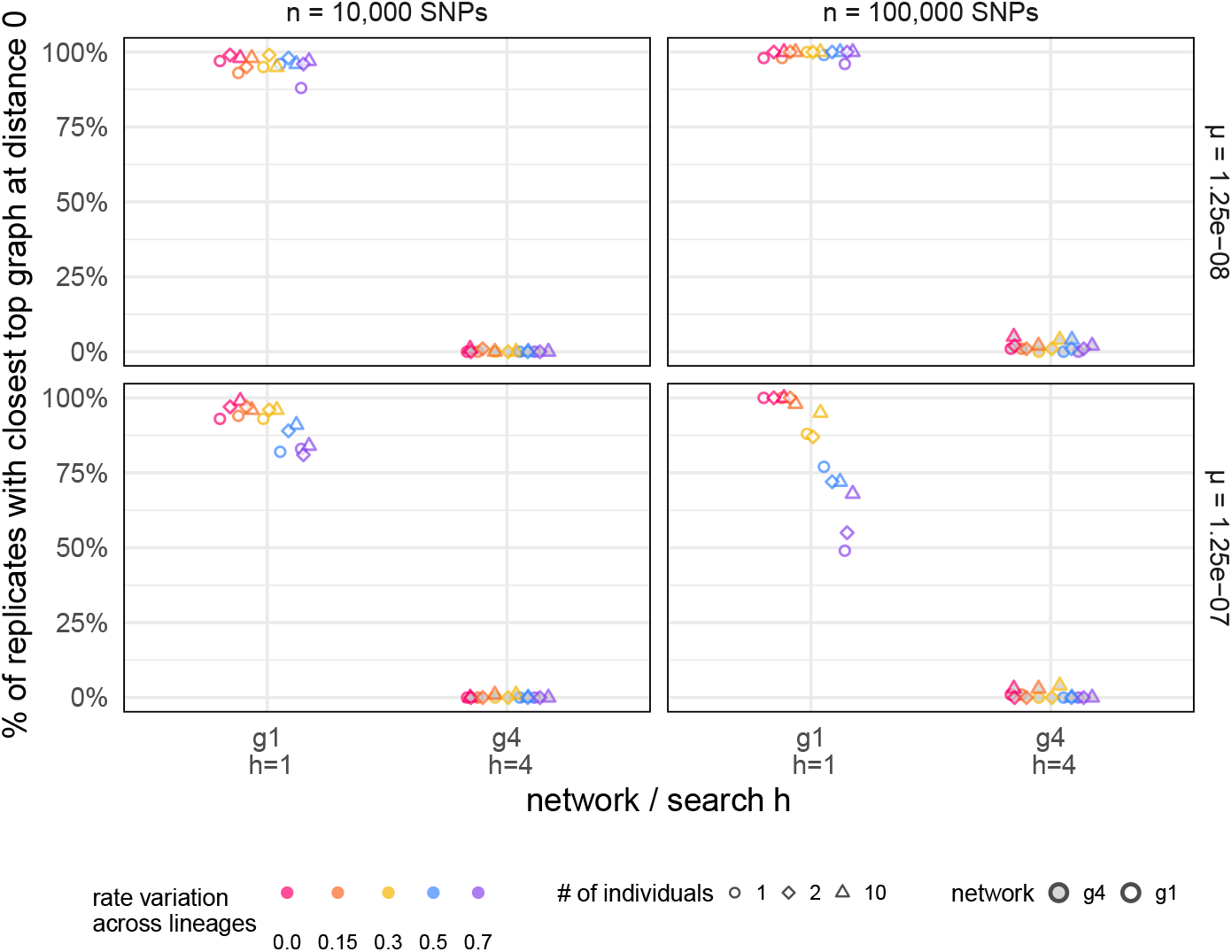
Percentage of data sets that have at least one top graph at hardwired-distance 0 from the true network `g1` or `g4` when searching for the true number of reticulations *h*, depending on mutation rate and data set size. Shape and color is as in Fig. S4.

Figs. S7 and S8 show the frequency with which the closest top graph was at distance 0 from the true network, given a search *h* equal to the true *h*. If *G* and g are trees or networks of level 1, then the hardwired-distance *d* is a proper distance: *G* = g exactly when *d*(*G*, `g`) = 0, and this criterion captures accuracy of network inference exactly. But for networks with *h* ≥ 2 reticulations and of level *ℓ* ≥ 2, if is possible that *d*(*G*, `g`) = 0 for *G* ≠ `g` (see examples in Fig. S3). So for our population networks `g` of level 2 or higher, including `g4`, the probability that at least top graph is at distance 0 from g is a relaxed (lenient) measure of accuracy. Still, even with this relaxation, `g4` was recovered with extremely low accuracy (at most 5%).

#### Distance of the closest top graph to the true network

Fig. 6 (for data generated under `g4`) and Fig. S9 (for data generated under `g1`) show the distribution of the hardwired-distance of the closest top graph to the true network, *d*_min_ = min {*d*(*G*, g) : *G* in top graph set} where `g` is the true network. To assess whether rate variation had an effect on this measure of accuracy, we conducted a Poisson regression on the transformed value *y* = *d*_min_*/*2 −*m* where *m* = 1 for `g` = `g1` if the search *h* = 0, *m* = 6 for `g` = `g4` if the search *h* = 0, *m* = 3 for `g` = `g4` if the search *h* = 1 and *m* = 0 otherwise (that is, if the search *h* is at least as large as the true *h*). This transformation was done to meet the Poisson distribution assumption for *y*, because *d*_min_ is a multiple of 2 (for resolved phylogenies), and because *d*_min_*/*2 is bounded by a minimum value when comparing phylogenies with different numbers of reticulation.

We used Poisson regression with our simulation parameters as predictors (all taken as factors with discrete levels): the true network, the search *h* (nested in network, as different *h* values were considered for `g1` and `g4`), the mutation rate, the number of biallelic sites, the level of rate variation across lineages, and no interaction effects. The Poisson regression model fit the data adequately (using both the residual deviance test and the Pearson chi-square test). There was overwhelming evidence for an effect of rate variation (likelihood ratio test statistic *X*^2^ = 241, *p <* 2 × 10^*−*16^). There was no discernible difference (*p* = 0.10) between a molecular clock (*σ* = 0) and the lowest level of rate variation considered here (*σ* = 0.15) but strong evidence (*p* = 0.001) of an increase in the average distance between *σ* = 0 and *σ* = 0.3 and overwhelming evidence for *σ* = 0.5 and 0.7 (*p <* 2 × 10^*−*16^), with increasingly more rate variation leading to an increasing distance between the closest top graph and the true network.

**Figure S9:**
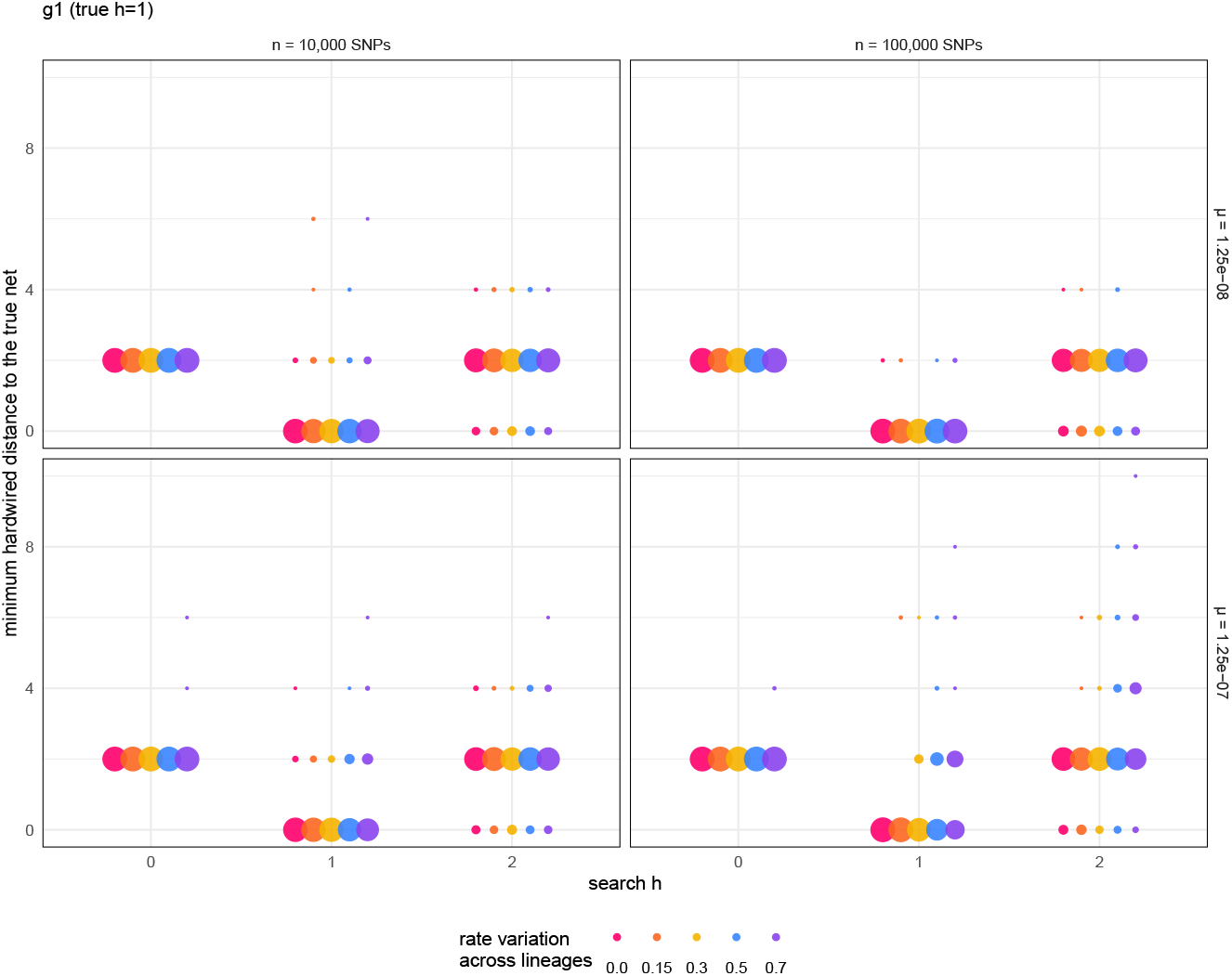
Distribution of the hardwired-distance between the true network and the closest top graph (that is, minimum hardwired-distance to the true network across top graphs), for data sets simulated under `g1`. The area of each circle is proportional to the percentage of replicate data sets, among data sets with a given combination of mutation rate *µ*, number of biallelic sites *n*, and maximum reticulation number *h* during graph search. Color denotes the amount of rate variation (the standard deviation, *σ* on a log-normal distribution with mean 1), with pink representing *σ* = 0 (a molecular clock), orange *σ* = 0.15, yellow *σ* = 0.3, blue *σ* = 0.5, and purple *σ* = 0.7.

### 6. Do top graphs display the true major tree?

We first show the probability that at least one top graph displays the true unrooted gene tree, for `g1` in Fig. S10 and `g4` in Fig. S11. Figs. S12 and S13 provide more details, showing the distance of the true major tree to the closest tree displayed in any topgraph.

**Figure S10:**
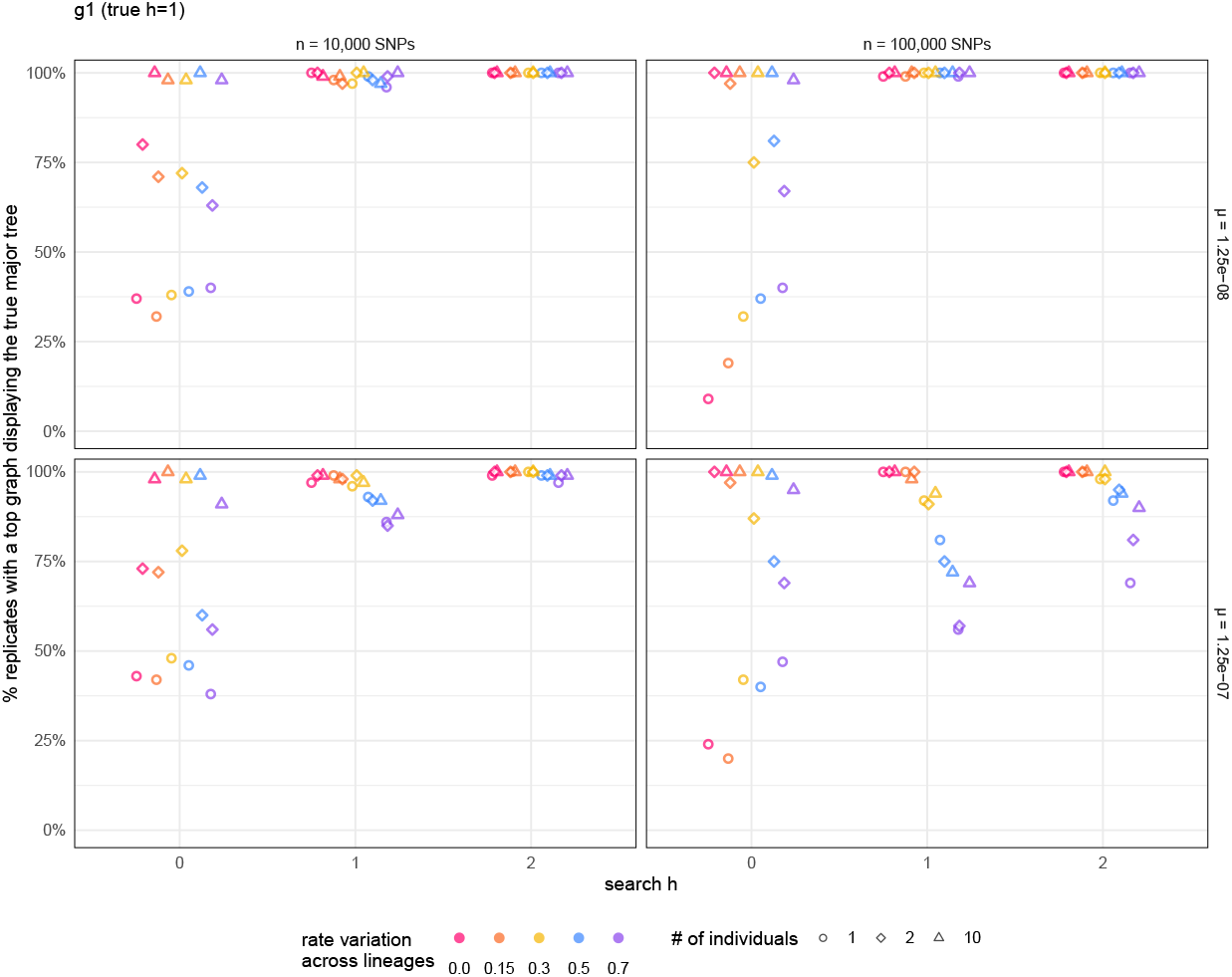
Percentage of data sets simulated under `g1` with at least one top graph displaying the true major tree, when searching for the true reticulation number *h*, depending on mutation rate *µ* and data set size. Shape and color is as in Fig. S4.

**Figure S11:**
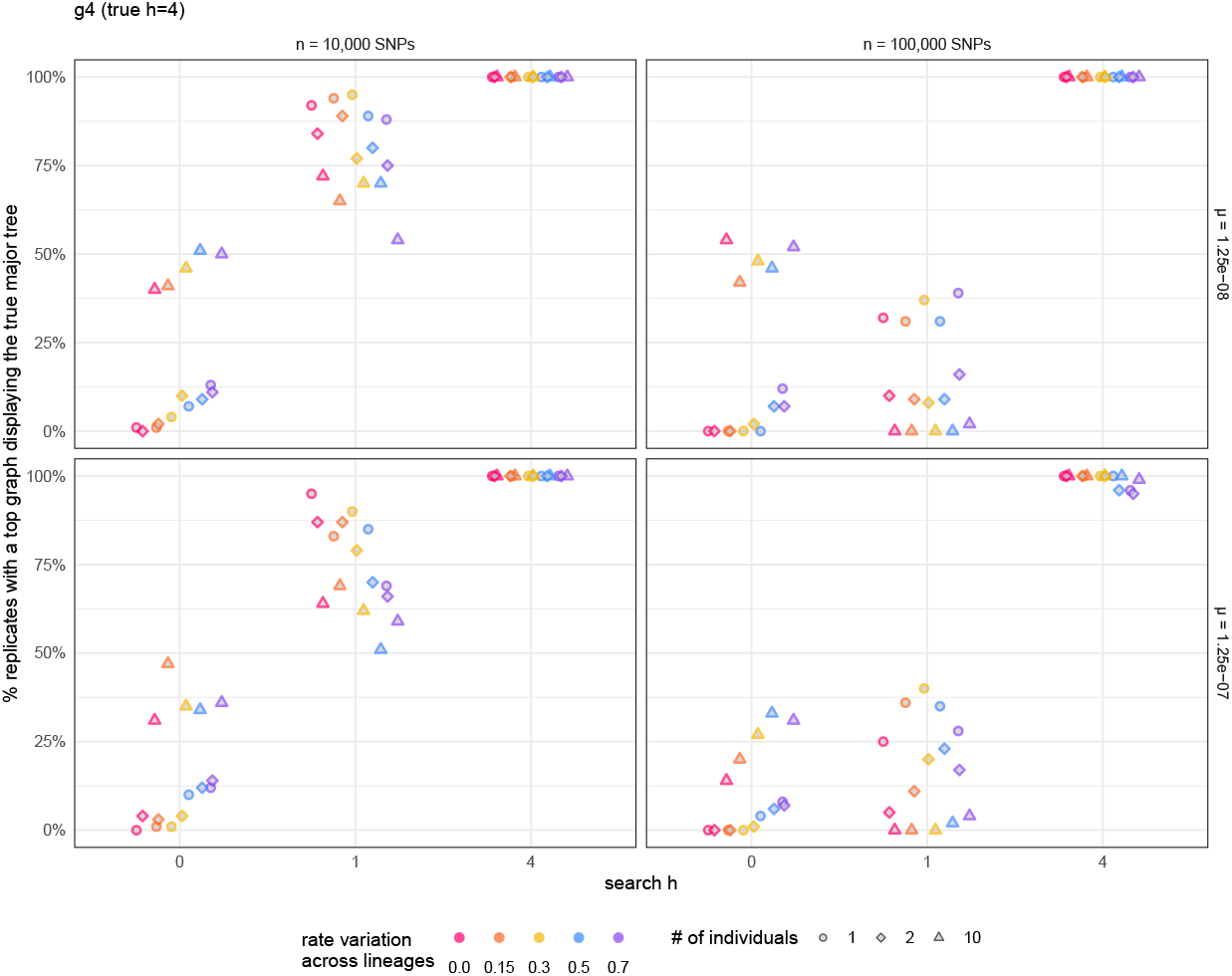
Percentage of data sets that have at least one top graph displaying the true major tree as in Fig. S10, but for data simulated under `g4`.

**Figure S12:**
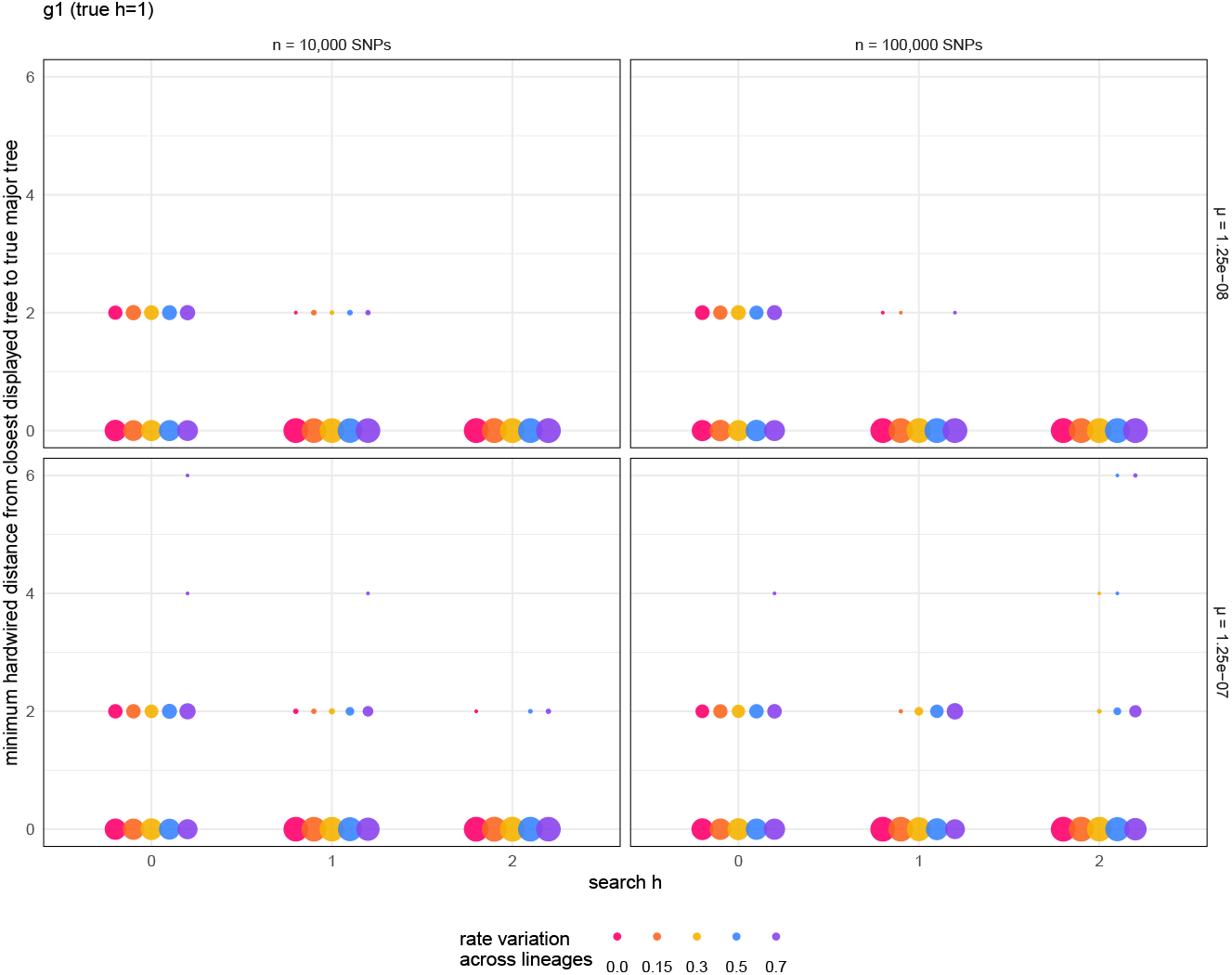
Distribution of the hardwired-distance between the true major tree and the closest top graphs’ displayed tree (that is, minimum hardwired-distance to the true major tree across displayed trees from top graphs), for data sets simulated under `g1`. The area of each circle is proportional to the percentage of replicate data sets, among the 300 data sets with a given combination of mutation rate *µ* and number of biallelic sites *n*. Color denotes the amount of rate variation as in Fig. S9.

**Figure S13:**
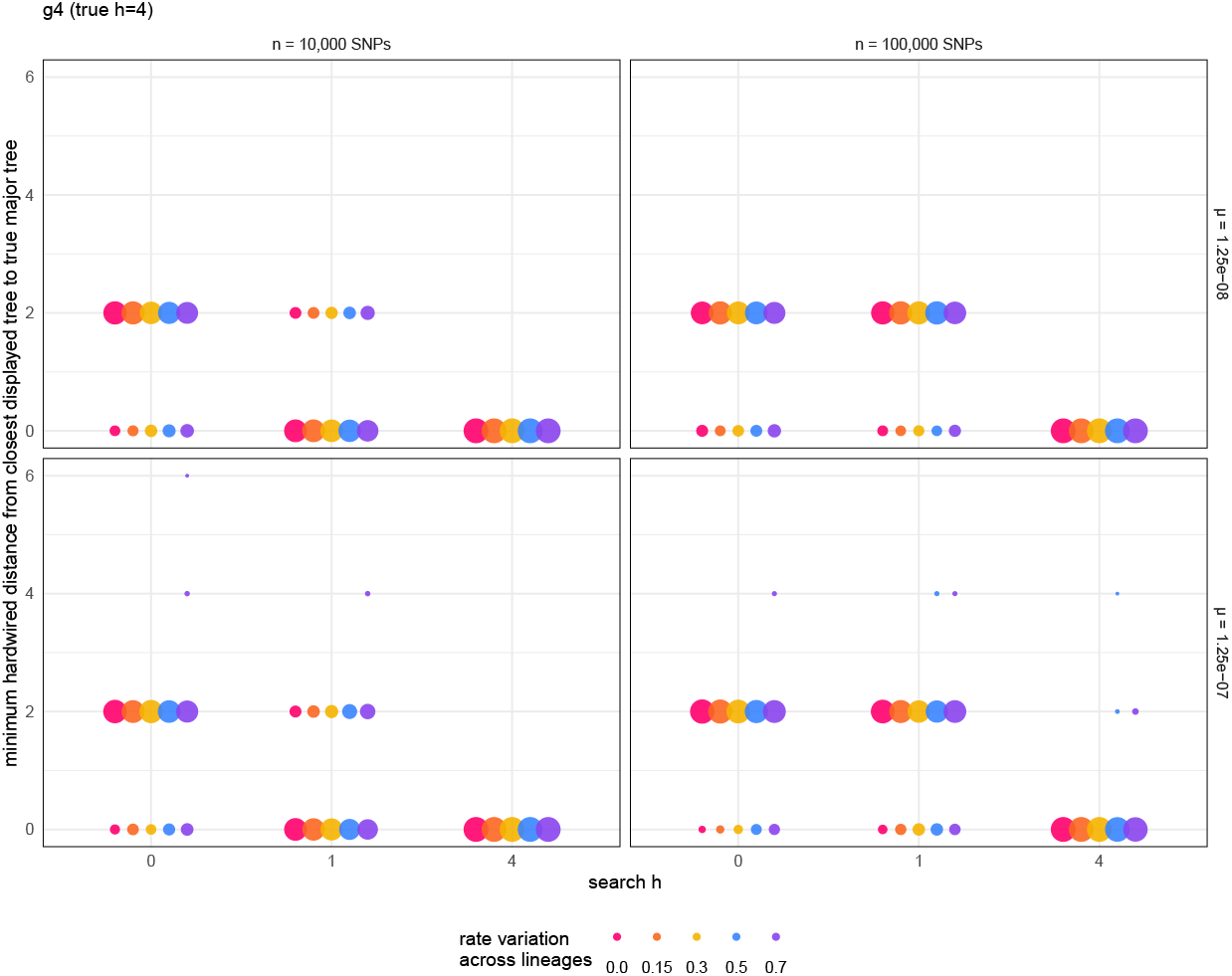
Hardwired-distance between the true major tree and the closest top graphs’ displayed tree as in Fig. S12, but for data simulated under `g4`.

### 7. Do top graphs display one or both near-major trees?

**Figure S14:**
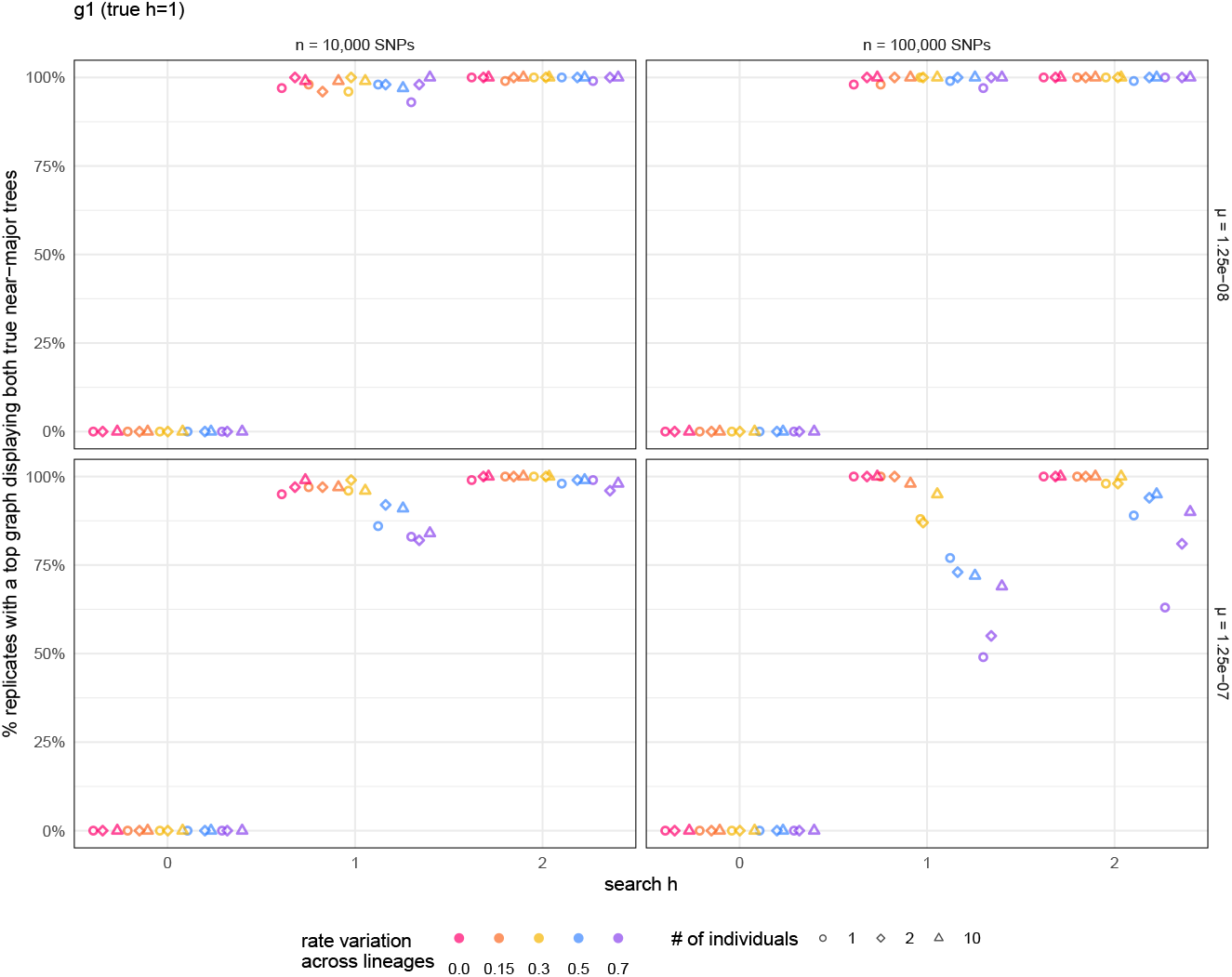
Percentage of 100 replicate data sets simulated under `g1` that have at least one top graph displaying both true near-major trees, depending on mutation rates *µ* and data set size. Top graphs were obtained by searching for high-likelihood graphs with at most “search h” reticulations. Shape and color is as in Fig. S4.

**Figure S15:**
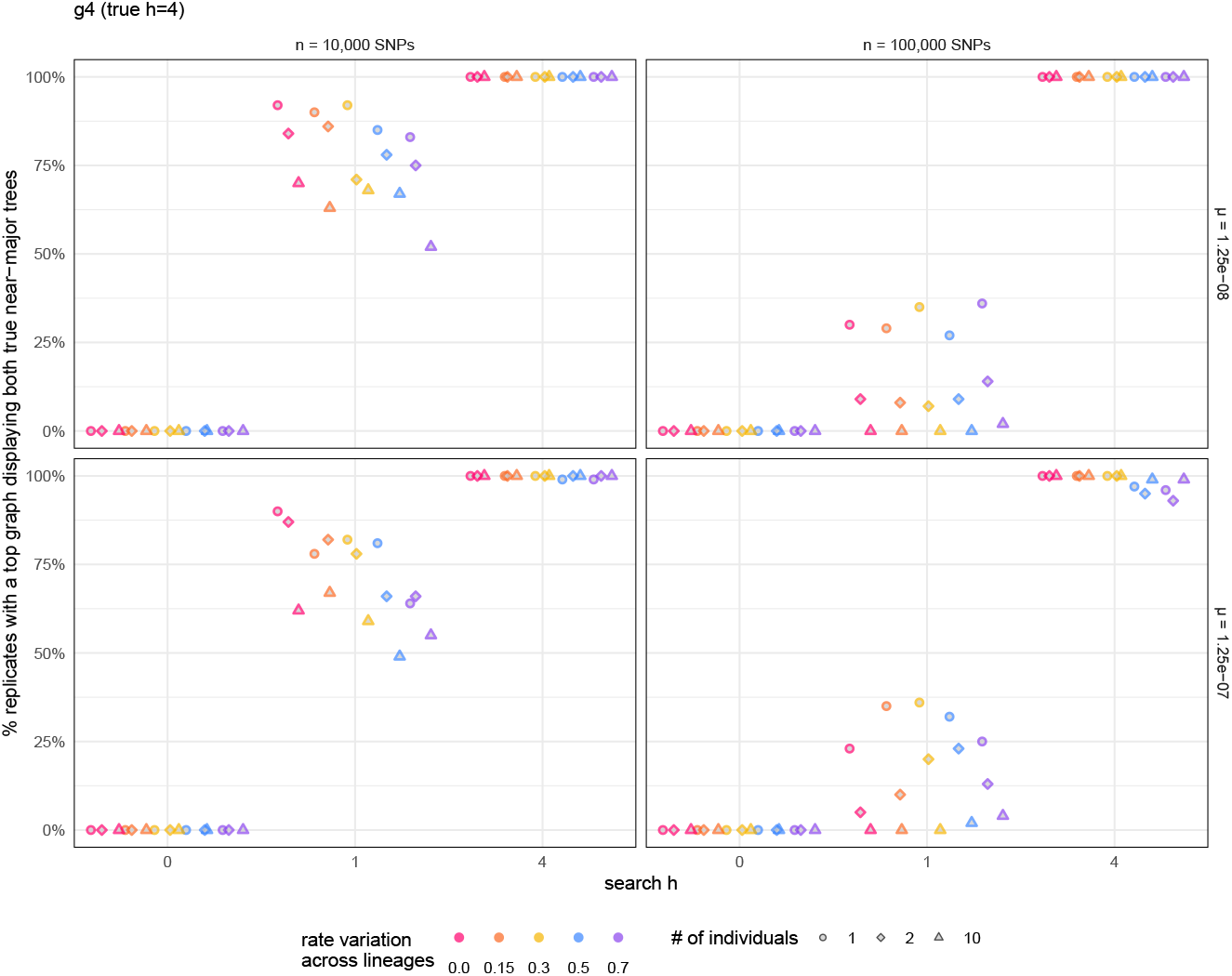
Percentage of data sets that have at least one top graph displaying both true near-major trees, as in Fig. S14, but for data sets simulated under `g4`.

**Figure S16:**
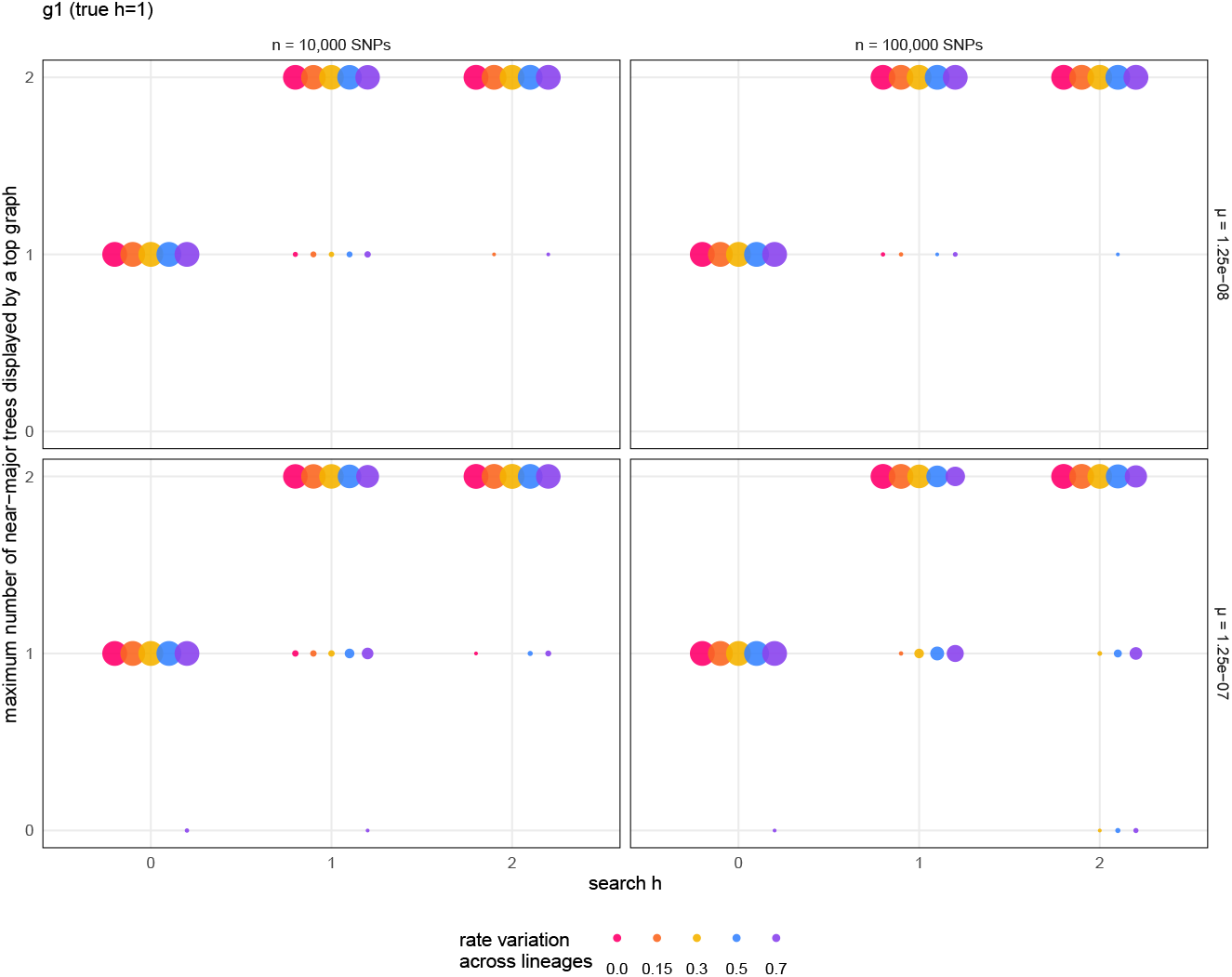
Maximum number of near-major trees displayed by a top graph (where the maximum is taken over all graphs in a replicate’s set of top graphs), for data sets simulated under `g1`. The area of each circle is proportional to the percentage of replicate data sets, among data sets with a given combination of mutation rate *µ*, number of biallelic sites *n*, and maximum reticulation number *h* during graph search. Color is as in Fig. S9.

**Figure S17:**
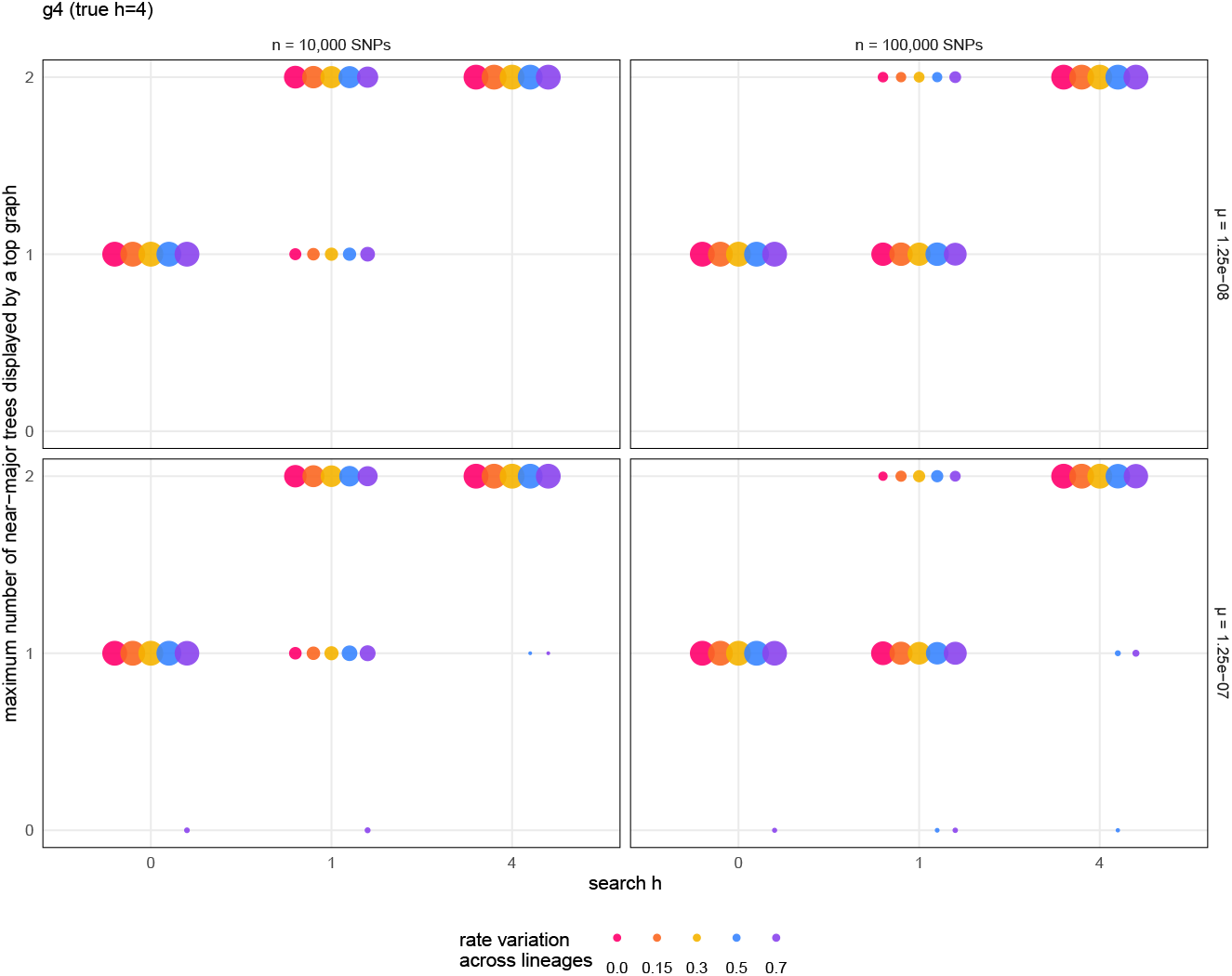
Maximum number of near-major trees displayed by a top graph (where the maximum is taken over all graphs in a replicate’s set of top graphs) for data sets simulated under `g4`. Area and color are as in Fig. S16.

### 8. How thoroughly was the graph space explored?

We might wonder if poor accuracy was due to a network having features that make it extremely difficult to recover, or if this was due to poor exploration of the graph space. To ensure that our graph search was sufficiently thorough, we increased search parameters (see Methods) within limits of running time constraints (see Supplementary section 4 and Table S1), and we compared the score of the best-scoring top graph *Ŝ* with the score of the true network *S*(g).

When searching for up to *h ≥* 1 reticulations under `g1`, or up to *h* = 4 reticulations under `g4`, we observed Δ*S* = *S*(g) *− Ŝ ≥* 0 all the time, as expected theoretically from a complete search, suggesting a sufficiently thorough search of the graph space.

**Figure S18:**
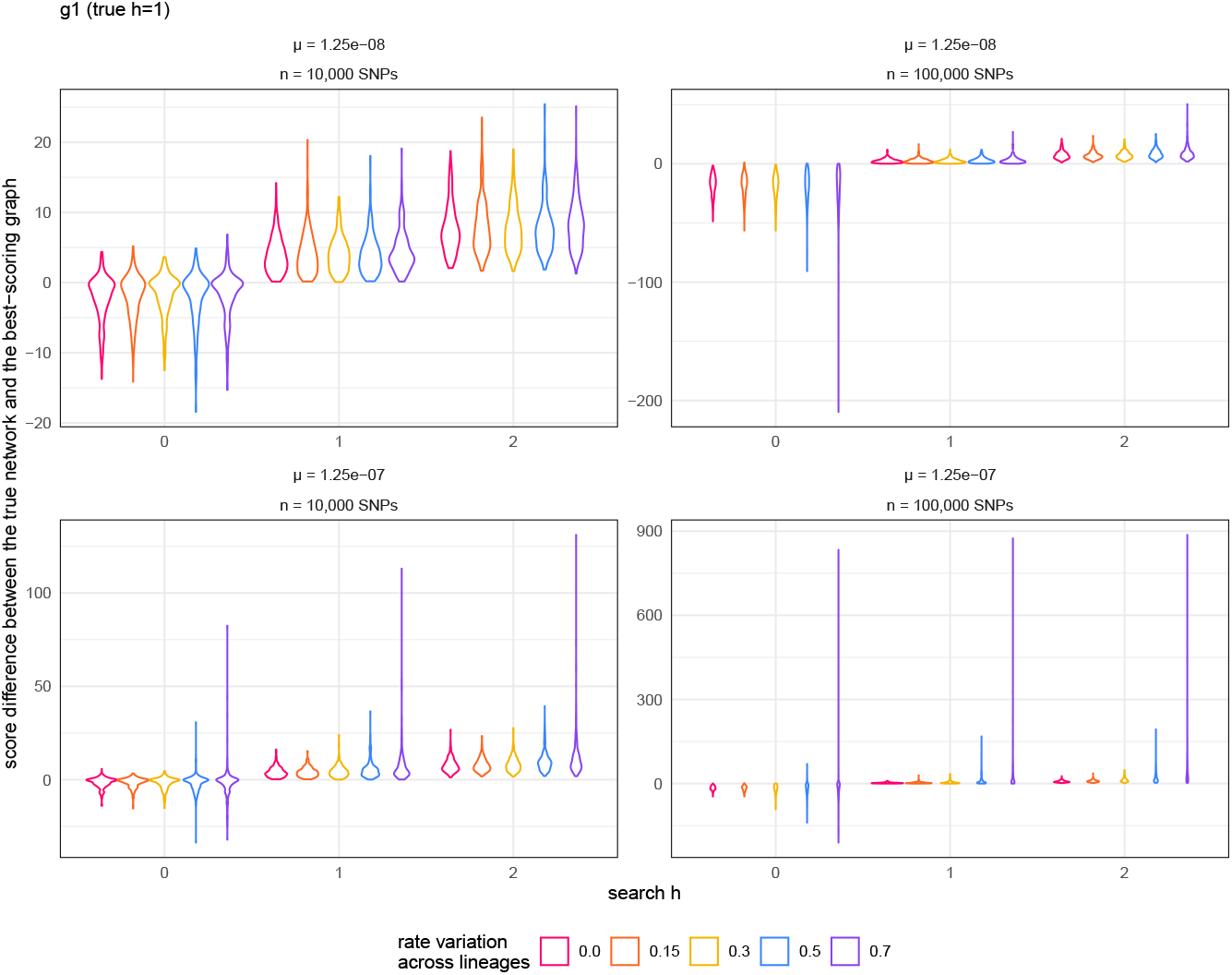
Distribution of the score difference between the true network and the best-scoring top graph, for data sets simulated under `g1`. Each violin represents the 300 replicate data sets with a given combination of mutation rate *µ* and number of biallelic sites *n*. Color denotes the amount of rate variation across lineages as in Fig. S4.

**Figure S19:**
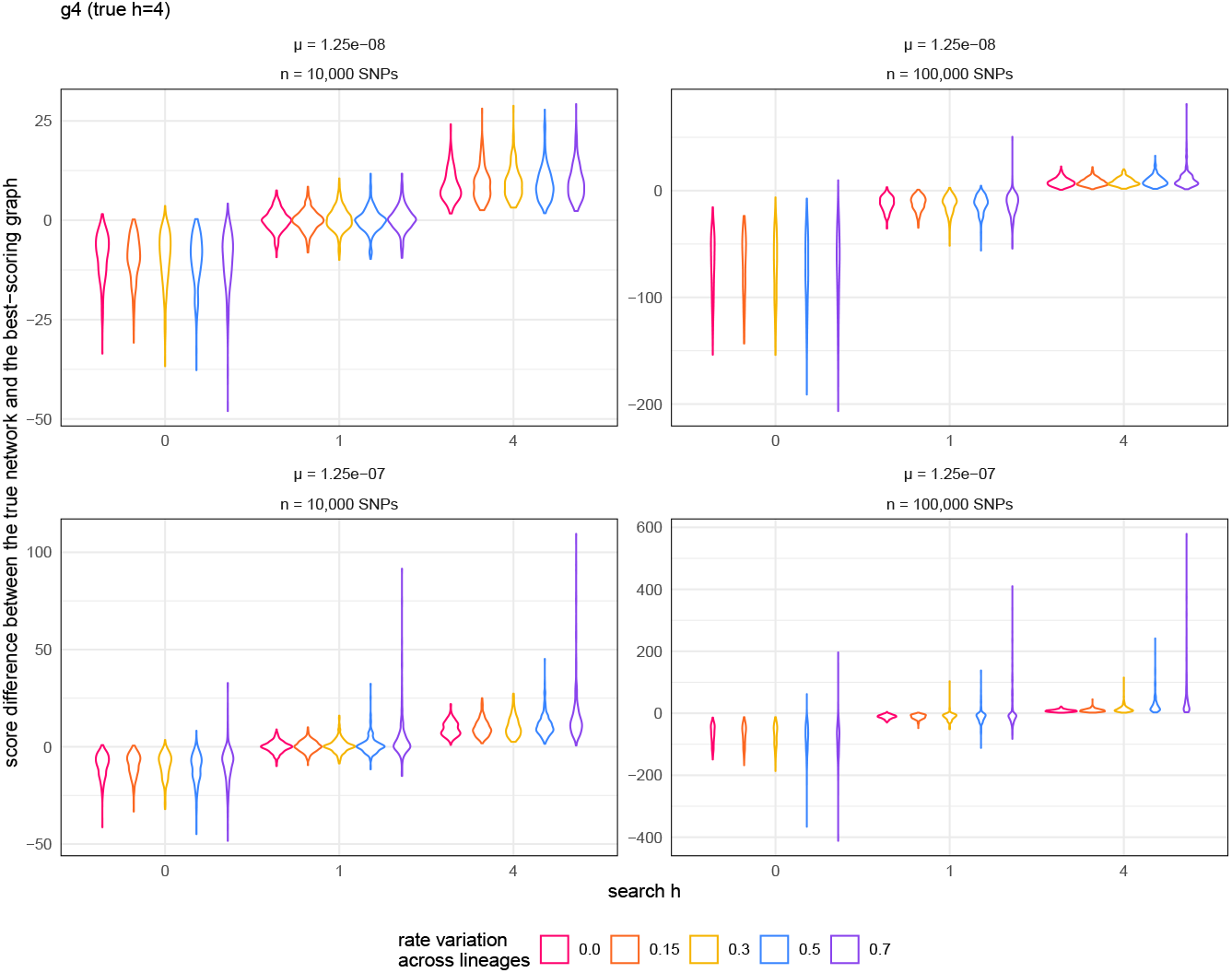
Score difference between the true network and the best-scoring graph as in Fig. S18, but for data simulated under `g4`.

### 9. How does graph filtering affect inference accuracy?

To obtain a list of “top graphs”, we filtered the list of all graphs returned by `find_graphs` across 50 independent runs, keeping the 5 best-scoring graphs and any graph scoring within 10 of the best score (*S > Ŝ* −10). This filtering strategy was quite conservative, as evidenced by the number of graphs retained and evaluated (see Table S2). Keeping a smaller list of top graphs is bound to reduce the accuracy of graph inference, unless the true network ranks high on the list of graphs sorted by their likelihood score. We report here the accuracy under three alternative filtering strategies, which consist in keeping: only the best-scoring graph (most liberal); the 5 highest scoring graphs; or the 5 highest scoring graphs and any graph scoring within 3 of the best score (*S > Ŝ −* 3).

#### Retaining only the highest scoring graph per replicate

There was no difference of accuracy when accuracy was measured by requiring at least one top graph to be ‘equal to’ versus ‘at hardwired-distance 0 of’ the true network. Therefore, only accuracy based on graph equality is shown below.

When evaluating only the best scoring graph, accuracy dropped sharply (Fig. S20 below). For networks without any hybridization with *c* = 3, accuracy dropped from over 98% under our conservative filtering to 51% at best, and to 0%-5% for 5 of these networks. These 5 networks were more complex that those recovered with higher accuracy: either with smaller cycles (4-cycles in all subnetworks), not tree-child, with 3 reticulations, or with low *γ* (g1-low*γ*, g2-l1-c44, g2-l2-t44, g2-l2-n54, g3-l3-c545). When there was one or two hybridizations with a minimum cycle size of *c* = 3, the best scoring graph was never the true graph.

**Figure S20:**
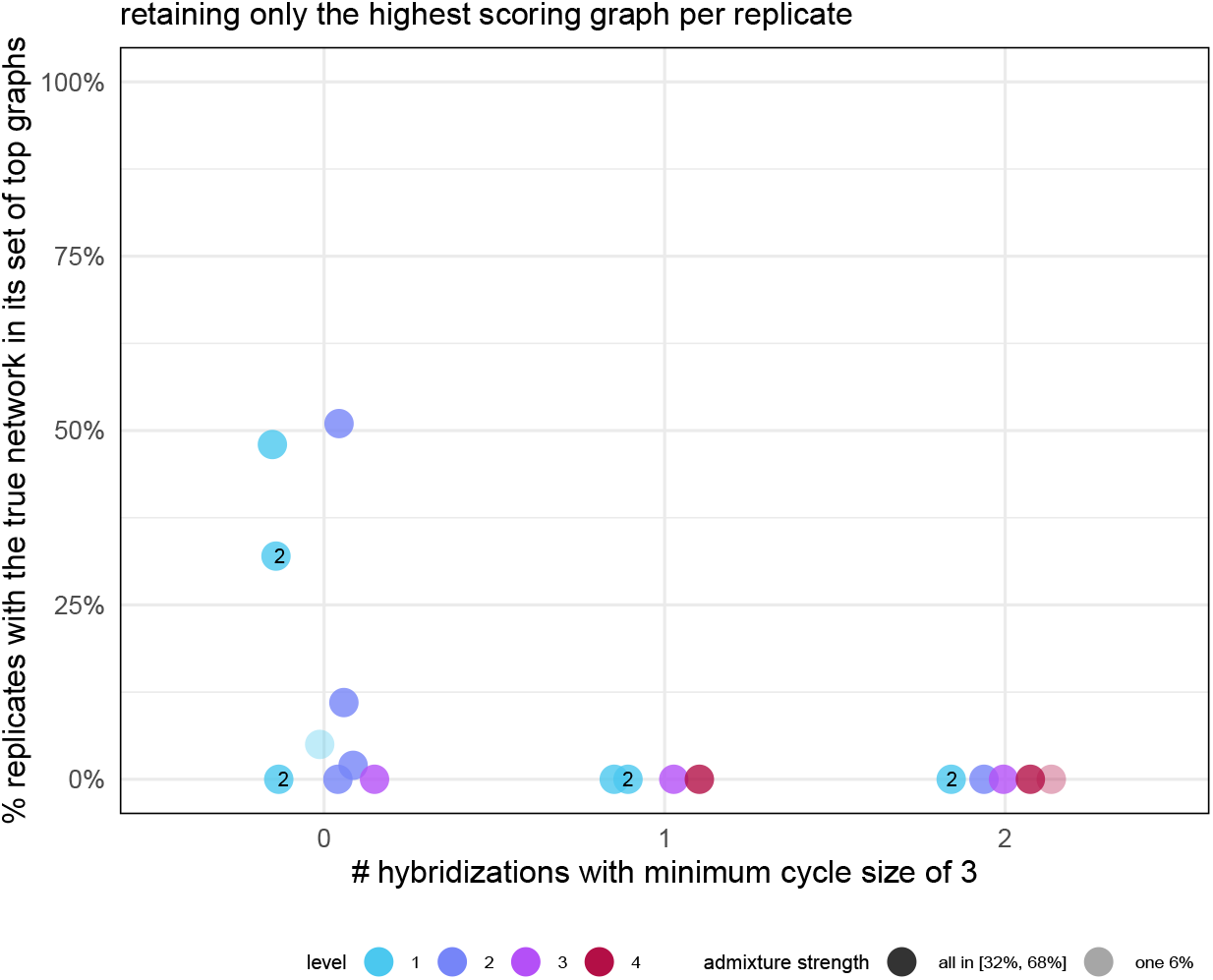
Percentage of data sets whose best scoring graph is the true network, when searching for the true number of reticulations *h* under baseline simulation parameters: 10 individuals per population, 100,000 biallelic sites, *µ* = 1.25 × 10^*−*8^ and a molecular clock. Points are labeled with the number of reticulations *h* if *h* does not match the level *ℓ*.

#### Retaining only the 5 highest scoring graphs per replicate

Again, we only show below accuracy based on graph equality, because accuracy was the same when requiring at least one top graph at hardwired-distance 0 of the true network.

For networks with one or two hybridizations with a minimum cycle size of *c* = 3, accuracy was again 0% when retaining the top 5 scoring graphs (Fig. S21). In cases without any hybridization with *c* = 3, the 5 networks of lowest (*≤* 5%) accuracy from 1 best-scoring graph still had low accuracy (0% to 26%) when keeping the top 5 scoring graphs. The other 4 networks were recovered with *≥* 85% accuracy (g1-high*γ*, g2-l1-c45, g2-l2-g46 and `g2-l2-t46`).

**Figure S21:**
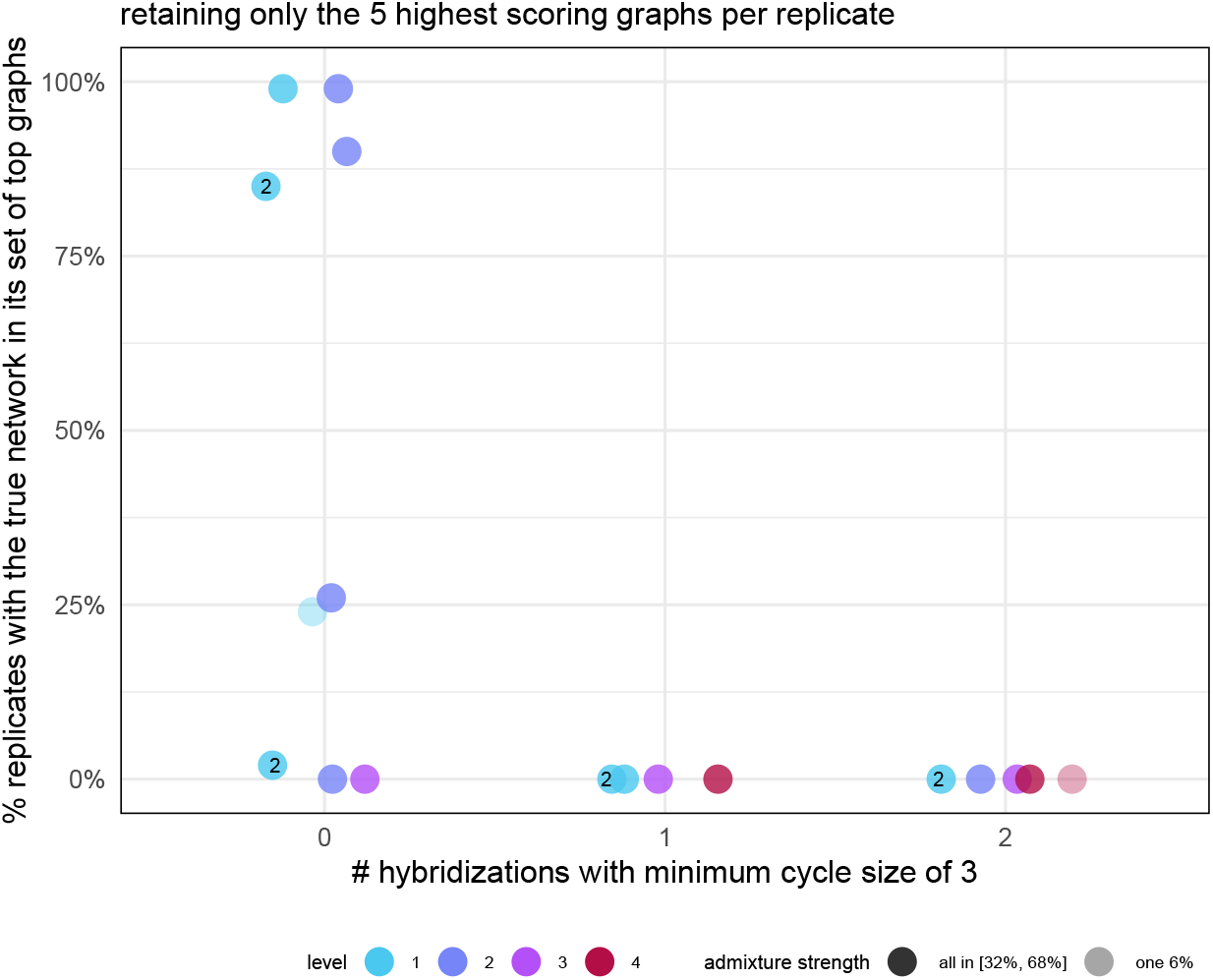
Percentage of data sets whose set of top graphs (5 best scoring graphs) contains the true network, when searching for the true number of reticulations *h* under baseline simulation parameters: 10 individuals per population, 100,000 biallelic sites, *µ* = 1.25 × 10^*−*8^ and a molecular clock. Points are labeled with the number of reticulations *h* if *h > ℓ*.

#### Retaining 5 highest scoring graphs and any graph scoring within 3 of the best score (*S > Ŝ* −3)

Compared to our baseline filtering, here we lower the score threshold for retaining graphs from 10 to 3. As noted in the discussion, a threshold drop of 3 in log-likelihood corresponds to an increase of 2 × 3 = 6 in the Bayesian information criterion (BIC) for graphs of the same *h* and same number of free edge parameters.

Obviously, keeping a larger set of top graphs (compared to only 5 graphs) improves accuracy (Figs. S22 and S23). But compared to a threshold of 10 in the log-likelihood score drop, accuracy dropped significantly. Therefore, this filtering strategy constitutes a trade-off between accuracy and size of the top graph set.

**Figure S22:**
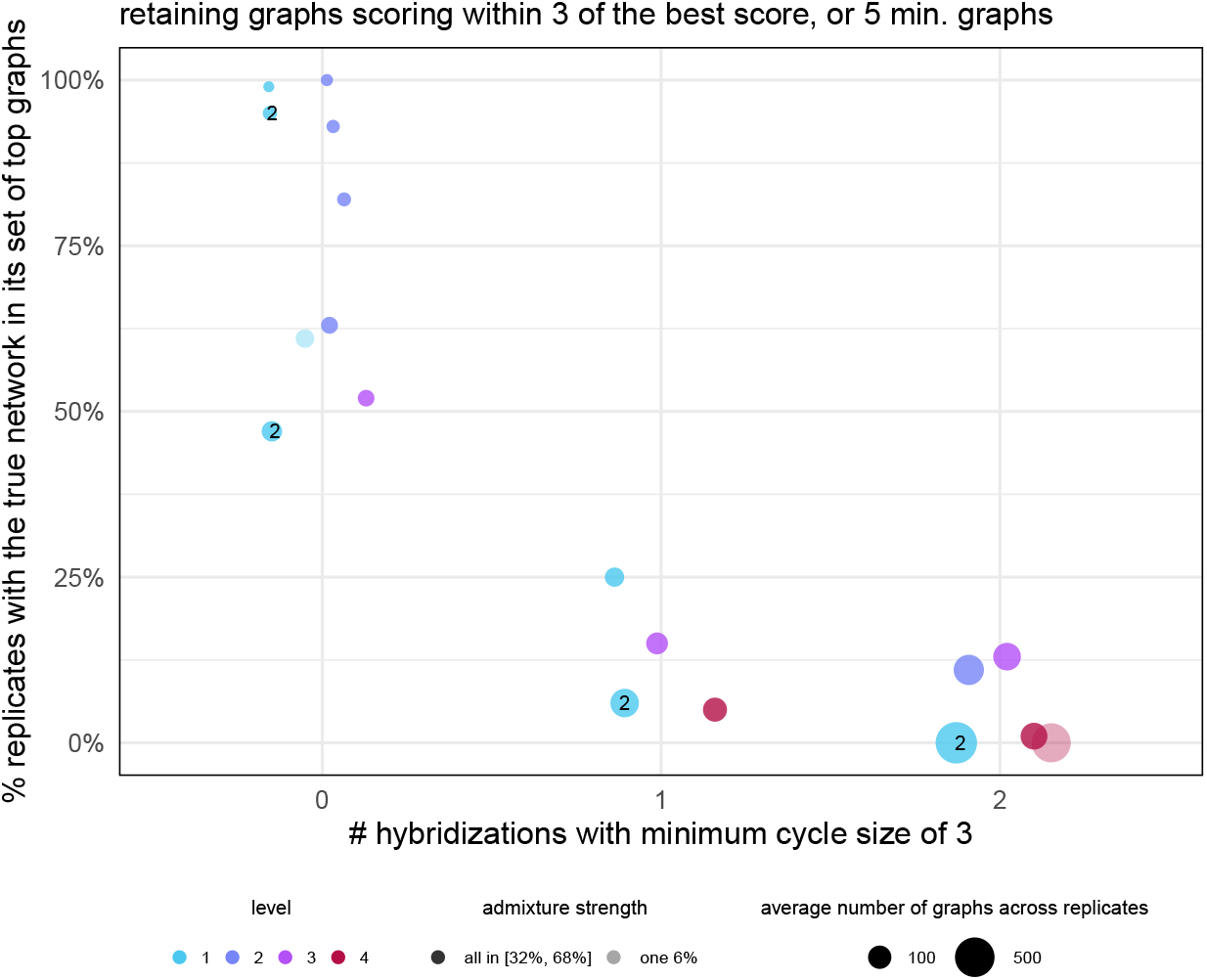
Percentage data sets whose set of top graphs (5 graphs and any graph scoring within 3 of the best score) contains the true network, when searching for the true number of reticulations *h* under baseline simulation parameters: 10 individuals per population, 100,000 biallelic sites, *µ* = 1.25 × 10^*−*8^ and a molecular clock. Points are labeled with the number of reticulations *h* if *h > ℓ*. Point area is proportional to the average number of top graphs.

**Figure S23:**
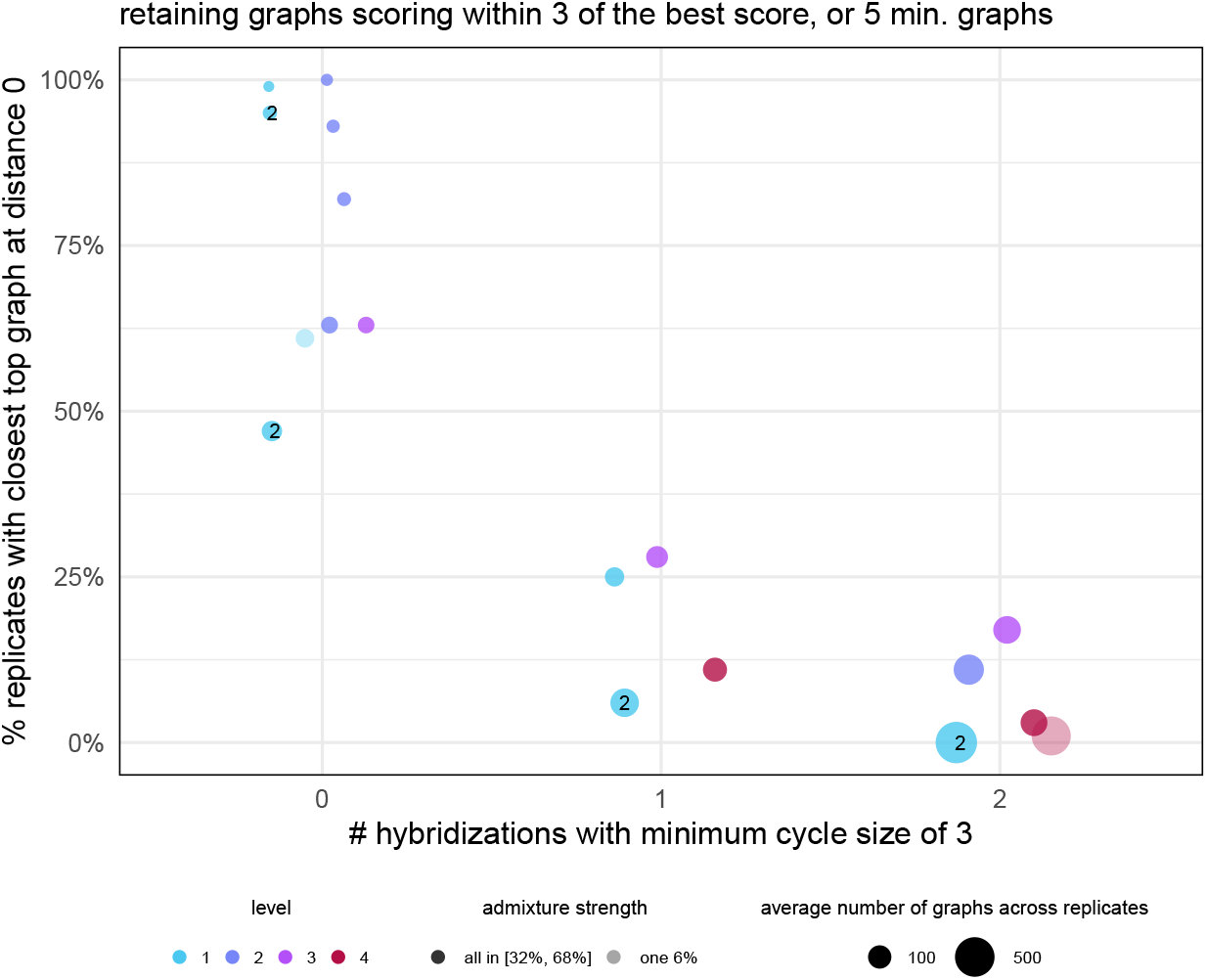
Percentage data sets whose set of top graphs (5 graphs and any graph scoring within 3 of the best score) contains at least one at hardwired-distance 0 from the true network, when searching for the true number of reticulations *h* under baseline simulation parameters: 10 individuals per population, 100,000 biallelic sites, *µ* = 1.25 × 10^*−*8^ and a molecular clock. Points are labeled with the number of reticulations *h* if *h > ℓ*. Point area is proportional to the average number of top graphs.

#### Assessing topological identifiability on g4

We extracted the 28 *f*_2_ values (from 28 pairs of populations) from the data simulated under `g4` and baseline parameters and averaged across replicates,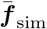. We then obtained the *f*_2_ values expected from `g4` under the infinite-site model used by admixtools, ***f*** _g4_, after estimating edge parameters on `g4` to fit 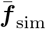. Fig. S24 (left) shows a perfect fit, with 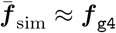. More specifically, the worst residual was 4.3 *×* 10^*−*5^, and the score of `g4` on 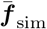, or sum of squares of residuals, was 7.7× 10^*−*9^. This excellent fit shows that using a simulation model different from the infinite-site model had little impact, under baseline parameters with low mutation rate *µ* and a molecular clock.

**Figure S24:**
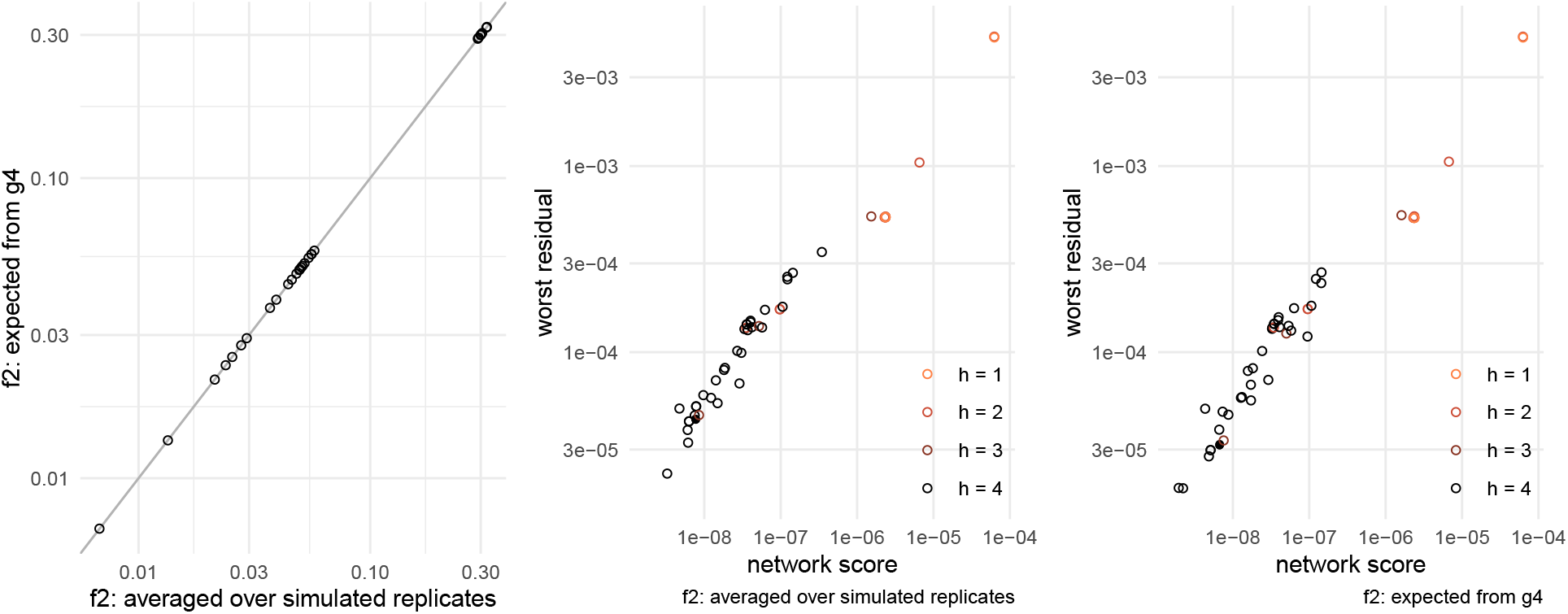
*f*_2_ values 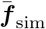 and ***f*** _g4_ (left); and model fit (score and worst residuals) of candidate networks fitted to 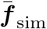(middle) or to ***f*** _g4_ (right), with `g4` indicated by a filled circle.

Fig. S24 shows the score and worst residual of candidate graphs, fitted either to 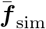 (middle) or to ***f*** _g4_ (right). Due to numerical precisions, `g4` (represented by a filled circle) does not have a perfect score of 0: *S*(`g4`) = 6.7 *×* 10^*−*9^ on ***f*** _g4_, only slightly better than 7.7 *×* 10^*−*9^ on 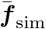. Other than `g4`, 12 topologies had a score *S <* 10^*−*8^ on 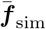, and 11 topologies had *S <* 10^*−*8^ on ***f*** _g4_. Since ***f*** _g4_ was calculated from `g4` itself, we used the score of `g4` as a measure of numerical precision, because *S*(`g4`) = 0 on ***f*** _g4_ theoretically. Fig. S25 shows `g4` and the 6 topologies that had *S*(`g`) *< S*(`g4`) under both ***f*** _g4_ and 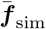. *F*-statistics expected from `g4` and from these 6 networks are identical, under admixtools’ model – hence a lack of identifiability.

**Figure S25:**
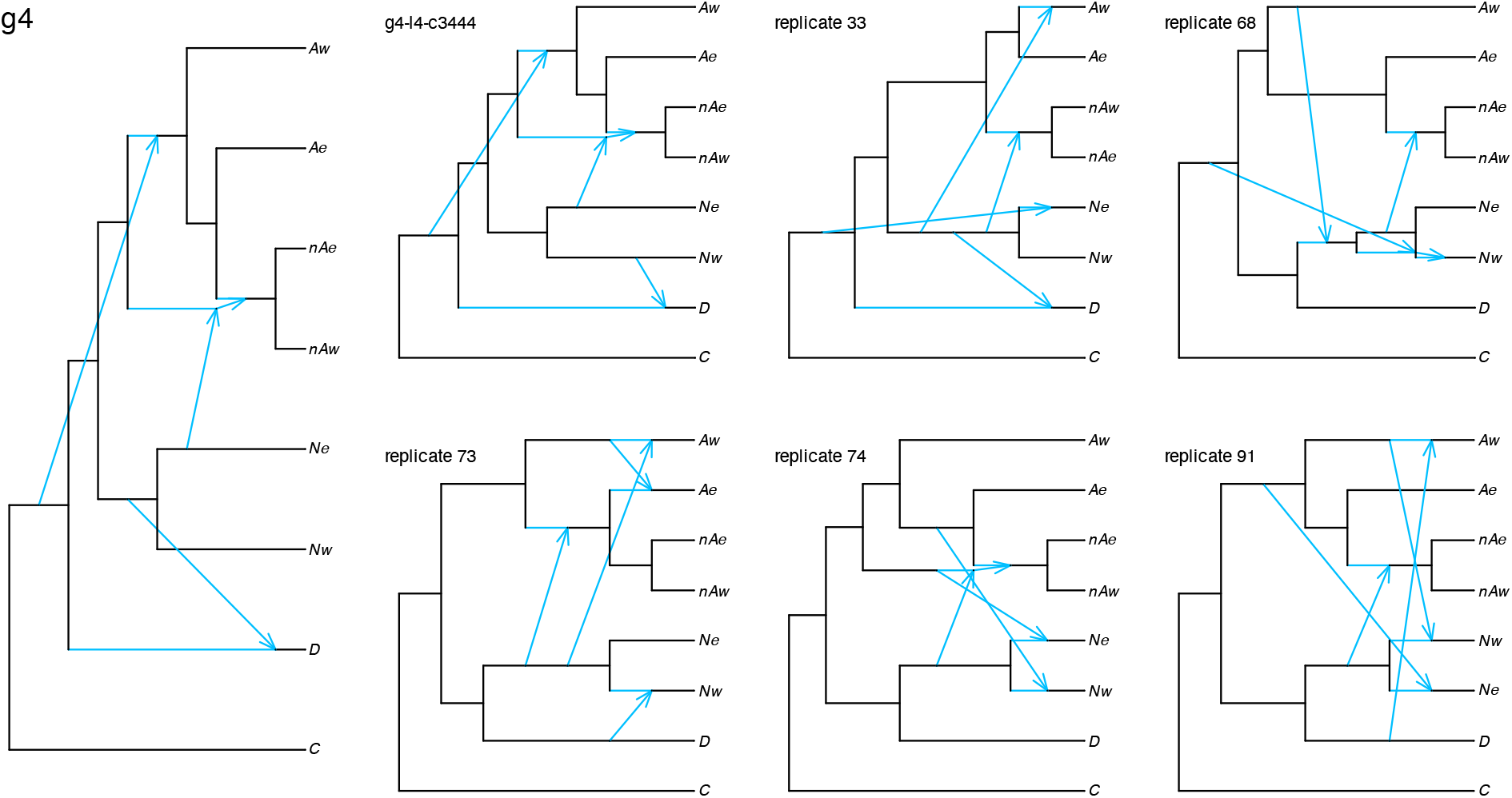
Network `g4` (left) and the 6 candidate networks fitting better than `g4` (*S*(g) *< S*(`g4`)) on both ***f***_g4_ and 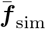. `g4`-l4-c3444 was used in our simulation (see Fig. S1, bottom row). Each other network was found as the highest-scoring graph from one replicate data set, whose index is indicated here. Population names are abbreviated are as in Fig. S1.

## References

H. Abu-Elmakarem, O. A. MacLean, F. Venter, L. J. Plenderleith, R. L. Culleton, B. H. Hahn, and P. M. Sharp. Remarkable evolutionary rate variations among lineages and among genome compartments in malaria parasites of mammals. Molecular Biology and Evolution, 41(12):msae243, 2024. doi:10.1093/molbev/msae243.

E. S. Allman, H. Baños, and J. A. Rhodes. Identifiability of species network topologies from genomic sequences using the logdet distance. Journal of Mathematical Biology, 84(5):35, 2022. doi:10.1007/s00285-022-01734-2.

E. S. Allman, H. Baños, J. D. Mitchell, and J. A. Rhodes. The tree of blobs of a species network: Identifiability under the coalescent. Journal of Mathematical Biology, 86(1):10, 2023. doi:10.1007/s00285-022-01838-9.

E. S. Allman, H. Baños, M. Garrote-Lopez, and J. A. Rhodes. Identifiability of level-1 species networks from gene tree quartets. Bulletin of Mathematical Biology, 86(110), 2024. doi:10.1007/s11538-024-01339-4.

E. S. Allman, C. Ané, H. Baños, and J. A. Rhodes. Beyond level-1: Identifiability of a class of galled tree-child networks. arXiv, 2025. doi:10.48550/arXiv.2504.21116.

L. Arbiza, M. Patricio, H. Dopazo, and D. Posada. Genome-wide heterogeneity of nucleotide substitution model fit. Genome biology and evolution, 3:896–908, 2011.

H. Baños. Identifying species network features from gene tree quartets under the coalescent model. Bulletin of Mathematical Biology, 81(2):494–534, 2019. doi:10.1007/s11538-018-0485-4.

L. A. Bergeron, S. Besenbacher, J. Zheng, P. Li, M. F. Bertelsen, B. Quintard, J. I. Hoffman, Z. Li, J. St. Leger, C. Shao, et al. Evolution of the germline mutation rate across vertebrates. Nature, 615(7951):285–291, 2023.

J. Bezanson, A. Edelman, S. Karpinski, and V. B. Shah. Julia: A fresh approach to numerical computing. SIAM Review, 59(1):65–98, 2017. doi:10.1137/141000671.

P. D. Blischak, J. Chifman, A. D. Wolfe, and L. S. Kubatko. HyDe: A python package for genome-scale hybridization detection. Systematic Biology, 67:821–829, 2018. doi:10.1093/sysbio/syy023.

M. Coll Macià, L. Skov, B. M. Peter, and M. H. Schierup. Different historical generation intervals in human populations inferred from neanderthal fragment lengths and mutation signatures. Nature Communications, 12(1):5317, 2021.

J. H. Degnan. Modeling hybridization under the network multispecies coalescent. Systematic Biology, 67(5):786–799, 2018. doi:10.1093/sysbio/syy040.

D. A. Duchêne, S. Duchêne, J. Stiller, R. Heller, and S. Y. W. Ho. ClockstaRX: Testing molecular clock hypotheses with genomic data. Genome Biology and Evolution, 16(4):evae064, 2024. doi:10.1093/gbe/evae064.

S. Duchêne, M. Molak, and S. Y. W. Ho. ClockstaR: choosing the number of relaxed-clock models in molecular phylogenetic analysis. Bioinformatics, 30(7):1017–1019, 2014. doi:10.1093/bioinformatics/btt665.

E. Y. Durand, N. Patterson, D. Reich, and M. Slatkin. Testing for ancient admixture between closely related populations. Molecular Biology and Evolution, 28:2239–2252, 2011. doi:10.1093/molbev/msr048.

A. K. Englander, M. Frohn, E. Gross, N. Holtgrefe, L. van Iersel, M. Jones, and S. Sullivant. Identifiability of phylogenetic level-2 networks under the Jukes-Cantor model. bioRxiv, 2025. doi:10.1101/2025.04.18.649493.

P. Flegontov, N.E. Altınışık, P. Changmai, N. Rohland, S. Mallick, N. Adamski, D. A. Bolnick, N. Broomandkhoshbacht, F. Candilio, B. J. Culleton, et al. Palaeo-eskimo genetic ancestry and the peopling of chukotka and north america. Nature, 570(7760):236–240, 2019.

P. Flegontov, U. Işıldak, R. Maier, E. Yüncü, P. Changmai, and D. Reich. Modeling of African population history using f-statistics can be highly biased and is not addressed by previously suggested SNP ascertainment schemes. bioRxiv, pages 2023–01, 2023.

T. Flouri, J. Huang, X. Jiao, P. Kapli, B. Rannala, and Z. Yang. Bayesian phylogenetic inference using relaxed-clocks and the multispecies coalescent. Molecular Biology and Evolution, 39(8):msac161, 2022. doi:10.1093/molbev/msac161.

J. Fogg and C. Ané. PhyloCoalSimulations v0.1.0. https://github.com/JuliaPhylo/PhyloCoalSimulations.jl, 2022. Last accessed: 2025-07-07.

J. Fogg, E. S. Allman, and C. Ané. PhyloCoalSimulations: a simulator for network multispecies coalescent models, including a new extension for the inheritance of gene flow. Systematic Biology, 72(5):1171–1179, 2023. doi:10.1093/sysbio/syad030.

L. E. Frankel and C. Ané. Summary Tests of Introgression Are Highly Sensitive to Rate Variation Across Lineages. Systematic Biology, page syad056, 09 2023. ISSN 1063-5157. doi:10.1093/sysbio/syad056. URL https://doi.org/10.1093/sysbio/syad056.

M. Gautier, R. Vitalis, L. Flori, and A. Estoup. f-statistics estimation and admixture graph construction with Pool-Seq or allele count data using the R package poolfstat. Molecular Ecology Resources, 22(4):1394–1416, 2022. doi:10.1111/1755-0998.13557.

X. Ge, Y. Lu, S. Chen, Y. Gao, L. Ma, L. Liu, J. Liu, X. Ma, L. Kang, and S. Xu. Genetic origins and adaptive evolution of the deng people on the tibetan plateau. Molecular Biology and Evolution, 40(10):msad205, 2023.

T. I. Gossmann, P. D. Keightley, and A. Eyre-Walker. The effect of variation in the effective population size on the rate of adaptive molecular evolution in eukaryotes. Genome biology and evolution, 4(5):658–667, 2012.

R. E. Green, J. Krause, A. W. Briggs, T. Maricic, U. Stenzel, M. Kircher, N. Patterson, H. Li, W. Zhai, M. H. Y. Fritz, N. F. Hansen, E. Y. Durand, A. S. Malaspinas, J. D. Jensen, T. Marques-Bonet, C. Alkan, K. Prüfer, M. Meyer, H. A. Burbano, J. M. Good, R. Schultz, A. Aximu-Petri, A. Butthof, B. Höber, B. Höffner, M. Siegemund, A. Weihmann, C. Nusbaum, E. S. Lander, C. Russ, N. Novod, J. Affourtit, M. Egholm, C. Verna, P. Rudan, D. Brajkovic, Željko Kucan, I. Gušic, V. B. Doronichev, L. V. Golovanova, C. Lalueza-Fox, M. D. L. Rasilla, J. Fortea, A. Rosas, R. W. Schmitz, P. L. Johnson, E. E. Eichler, D. Falush, E. Birney, J. C. Mullikin, M. Slatkin, R. Nielsen, J. Kelso, M. Lachmann, D. Reich, and S. Pääbo. A draft sequence of the Neandertal genome. Science, 328:710–722, 5 2010. doi:10.1126/science.1188021.

E. Gross, L. van Iersel, R. Janssen, M. Jones, C. Long, and Y. Murakami. Distinguishing level-1 phylogenetic networks on the basis of data generated by Markov processes. Journal of Mathematical Biology, 83:32, 2021. doi:10.1007/s00285-021-01653-8.

R. M. Gutaker, S. C. Groen, E. S. Bellis, J. Y. Choi, I. S. Pires, R. K. Bocinsky, E. R. Slayton, O. Wilkins, C. C. Castillo, S. Negrão, et al. Genomic history and ecology of the geographic spread of rice. Nature Plants, 6(5):492–502, 2020.

W. Haak, I. Lazaridis, N. Patterson, N. Rohland, S. Mallick, B. Llamas, G. Brandt, S. Nordenfelt, E. Harney, K. Stewardson, et al. Massive migration from the steppe was a source for indo-european languages in europe. Nature, 522 (7555):207–211, 2015.

M. W. Hahn and M. S. Hibbins. A three-sample test for introgression. Molecular Biology and Evolution, 36:2878–2882, 2019. doi:10.1093/molbev/msz178.

É. Harney, N. Patterson, D. Reich, and J. Wakeley. Assessing the performance of qpadm: a statistical tool for studying population admixture. Genetics, 217(4):iyaa045, 2021.

K. T. Huber, V. Moulton, and A. Spillner. Computing consensus networks for collections of 1-nested phylogenetic networks. Journal of Graph Algorithms and Applications, 27(7):541–563, 2023.

D. H. Huson, R. Rupp, and C. Scornavacca. Phylogenetic Networks: Concepts, Algorithms and Applications. Cambridge University Press, Cambridge, 2010.

J. Kamm, J. Terhorst, R. Durbin, and Y. S. Song. Efficiently inferring the demographic history of many populations with allele count data. Journal of the American Statistical Association, 115(531):1472–1487, 2020.

S. Kong, D. L. Swofford, and L. S. Kubatko. Inference of phylogenetic networks from sequence data using composite likelihood. Systematic Biology, 74(1):53–69, 2025. doi:10.1093/sysbio/syae054.

T. Koppetsch, M. Malinsky, and M. Matschiner. Towards reliable detection of introgression in the presence of among-species rate variation. Systematic biology, 73(5):769–788, 2024.

I. Lazaridis, N. Patterson, A. Mittnik, G. Renaud, S. Mallick, K. Kirsanow, P. H. Sudmant, J. G. Schraiber, S. Castellano, M. Lipson, B. Berger, C. Economou, R. Bollongino, Q. Fu, K. I. Bos, S. Nordenfelt, H. Li, C. de Filippo, K. Prüfer, S. Sawyer, C. Posth, W. Haak, F. Hallgren, E. Fornander, N. Rohland, D. Delsate, M. Francken, J.-M. Guinet, J. Wahl, G. Ayodo, H. A. Babiker, G. Bailliet, E. Balanovska, O. Balanovsky, R. Barrantes, G. Bedoya, H. Ben-Ami, J. Bene, F. Berrada, C. M. Bravi, F. Brisighelli, G. B. J. Busby, F. Cali, M. Churnosov, D. E. C. Cole, D. Corach, L. Damba, G. van Driem, S. Dryomov, J.-M. Dugoujon, S. A. Fedorova, I. Gallego Romero, M. Gubina, M. Hammer, B. M. Henn, T. Hervig, U. Hodoglugil, A. R. Jha, S. Karachanak-Yankova, R. Khusainova, E. Khusnutdinova, R. Kittles, T. Kivisild, W. Klitz, V. Kučinskas, A. Kushniarevich, L. Laredj, S. Litvinov, T. Loukidis, R. W. Mahley, B. Melegh, E. Metspalu, J. Molina, J. Mountain, K. Näkkäläjärvi, D. Nesheva, T. Nyambo, L. Osipova, J. Parik, F. Platonov, O. Posukh, V. Romano, F. Rothhammer, I. Rudan, R. Ruizbakiev, H. Sahakyan, A. Sajantila, A. Salas, E. B. Starikovskaya, A. Tarekegn, D. Toncheva, S. Turdikulova, I. Uktveryte, O. Utevska, R. Vasquez, M. Villena, M. Voevoda, C. A. Winkler, L. Yepiskoposyan, P. Zalloua, T. Zemunik, A. Cooper, C. Capelli, M. G. Thomas, A. Ruiz-Linares, S. A. Tishkoff, L. Singh, K. Thangaraj, R. Villems, D. Comas, R. Sukernik, M. Metspalu, M. Meyer, E. E. Eichler, J. Burger, M. Slatkin, S. Pääbo, J. Kelso, D. Reich, and J. Krause. Ancient human genomes suggest three ancestral populations for present-day Europeans. Nature, 513(7518):409–413, 2014. doi:10.1038/nature13673.

J. Lehtonen and R. Lanfear. Generation time, life history and the substitution rate of neutral mutations. Biology letters, 10(11):20140801, 2014.

P. Librado, N. Khan, A. Fages, M. A. Kusliy, T. Suchan, L. Tonasso-Calvière, S. Schiavinato, D. Alioglu, A. Fromentier, A. Perdereau, et al. The origins and spread of domestic horses from the western eurasian steppes. Nature, 598(7882): 634–640, 2021.

M. Lipson. Applying f4-statistics and admixture graphs: Theory and examples. Molecular Ecology Resources, 20(6):1658–1667, 2020. doi:10.1111/1755-0998.13230.

M. Lipson, O. Cheronet, S. Mallick, N. Rohland, M. Oxenham, M. Pietrusewsky, T. O. Pryce, A. Willis, H. Matsumura, H. Buckley, et al. Ancient genomes document multiple waves of migration in southeast asian prehistory. Science, 361(6397):92–95, 2018.

R. Maier, P. Flegontov, O. Flegontova, U. Isildak, P. Changmai, and D. Reich. On the limits of fitting complex models of population history to f-statistics. Elife, 12:e85492, 2023.

M. Malinsky, M. Matschiner, and H. Svardal. Dsuite-fast d-statistics and related admixture evidence from vcf files. Molecular ecology resources, 21(2):584–595, 2021.

A. Massimo, Le Trang, C. Corrado, B. Alessandra, and C. June. openalexr: An r-tool for collecting bibliometric data from openalex. The R Journal, 15:167–180, 2024. ISSN 2073-4859. doi:10.32614/RJ-2023-089.

J. V. Moreno-Mayar, B. A. Potter, L. Vinner, M. Steinrücken, S. Rasmussen, J. Terhorst, J. A. Kamm, A. Albrechtsen, A.-S. Malaspinas, M. Sikora, et al. Terminal pleistocene alaskan genome reveals first founding population of native americans. Nature, 553(7687):203–207, 2018.

M. Narasimhan, N. Patterson, P. Moorjani, N. Rohland, R. Bernardos, S. Mallick, I. Lazaridis, N. Nakatsuka, I. Olalde, M. Lipson, et al. The formation of human populations in south and central asia. Science, 365(6457): eaat7487, 2019.

S. V. Nielsen, A. H. Vaughn, K. Leppälä, M. J. Landis, T. Mailund, and R. Nielsen. Bayesian inference of admixture graphs on native american and arctic populations. PLOS Genetics, 19(2):1–22, 2023. doi:10.1371/journal.pgen.1010410.

N. Patterson, P. Moorjani, Y. Luo, S. Mallick, N. Rohland, Y. Zhan, T. Genschoreck, T. Webster, and D. Reich. Ancient admixture in human history. Genetics, 192(3):1065–1093, 2012.

K. M. Peck and A. S. Lauring. Complexities of viral mutation rates. Journal of virology, 92(14):10–1128, 2018.

B. M. Peter. Admixture, population structure, and F-statistics. Genetics, 202(4):1485–1501, 2016. doi:10.1534/genetics.115.183913.

B. M. Peter. A geometric relationship of f2, f3 and f4-statistics with principal component analysis. Philosophical Transactions of the Royal Society B: Biological Sciences, 377(1852):20200413, 2022. doi:10.1098/rstb.2020.0413.

N. Petit and A. Barbadilla. Selection efficiency and effective population size in drosophila species. Journal of Evolutionary Biology, 22(3):515–526, 2009.

J. Priem, H. Piwowar, and R. Orr. Openalex: A fully-open index of scholarly works, authors, venues, institutions, and concepts, 2022. URL https://arxiv.org/abs/2205.01833.

A. E. Raftery. Bayesian model selection in social research. Sociological Methodology, 25:111–163, 1995.

A. P. Ragsdale and S. Gravel. Models of archaic admixture and recent history from two-locus statistics. PLOS Genetics, 15(6):1–19, 2019. doi:10.1371/journal.pgen.1008204.

A. P. Ragsdale, T. D. Weaver, E. G. Atkinson, E. G. Hoal, M. Möller, B. M. Henn, and S. Gravel. A weakly structured stem for human origins in Africa. Nature, 617(7962):755–763, 2023a. doi:10.1038/s41586-023-06055-y.

A. P. Ragsdale, T. D. Weaver, E. G. Atkinson, E. G. Hoal, M. Möller, B. M. Henn, and S. Gravel. A weakly structured stem for human origins in africa. Nature, 617(7962):755–763, 2023b.

A. Rambaut and N. C. Grass. Seq-Gen: an application for the Monte Carlo simulation of DNA sequence evolution along phylogenetic trees. Bioinformatics, 13(3):235–238, 1997.

J. A. Rhodes, H. Baños, J. Xu, and C. Ané. Identifying circular orders for blobs in phylogenetic networks. Advances in Applied Mathematics, 163:102804, 2025. doi:10.1016/j.aam.2024.102804.

A. R. Rogers. Legofit: estimating population history from genetic data. BMC bioinformatics, 20:1–10, 2019.

A. Scally and R. Durbin. Revising the human mutation rate: implications for understanding human evolution. Nature Reviews Genetics, 13(10):745–753, 2012.

S. A. Smith and M. J. Donoghue. Rates of molecular evolution are linked to life history in flowering plants. science, 322(5898):86–89, 2008.

S. Snir, Y. I. Wolf, and E. V. Koonin. Universal pacemaker of genome evolution. PLOS Computational Biology, 8(11):1–9, 2012. doi:10.1371/journal.pcbi.1002785.

C. Solís-Lemus and C. Ané. Inferring phylogenetic networks with maximum pseudolikelihood under incomplete lineage sorting. PLoS Genetics, 12(3):e1005896, 2016. doi:10.1371/journal.pgen.1005896.

S. Soraggi and C. Wiuf. General theory for stochastic admixture graphs and f-statistics. Theoretical Population Biology, 125:56–66, 2019. doi:10.1016/j.tpb.2018.12.002.

Y. Thawornwattana, J. Huang, T. Flouri, J. Mallet, and Z. Yang. Inferring the direction of introgression using genomic sequence data. Molecular Biology and Evolution, 40(8):msad178, 2023. doi:10.1093/molbev/msad178.

J. A. Thomas, J. J. Welch, R. Lanfear, and L. Bromham. A generation time effect on the rate of molecular evolution in invertebrates. Molecular biology and evolution, 27(5):1173–1180, 2010.

T. van der Valk, P. Pečnerová, D. Díez-del Molino, A. Bergström, J. Oppenheimer, S. Hartmann, G. Xenikoudakis, J. A. Thomas, M. Dehasque, E. Sağlıcan, et al. Million-year-old DNA sheds light on the genomic history of mammoths. Nature, 591(7849):265–269, 2021.

C.-C. Wang, H.-Y. Yeh, A. N. Popov, H.-Q. Zhang, H. Matsumura, K. Sirak, O. Cheronet, A. Kovalev, N. Rohland, A. M. Kim, et al. Genomic insights into the formation of human populations in east asia. Nature, 591(7850):413–419, 2021.

R. J. Wang, S. I. Al-Saffar, J. Rogers, and M. W. Hahn. Human generation times across the past 250,000 years. Science Advances, 9(1):eabm7047, 2023.

Y. Wang and D. J. Obbard. Experimental estimates of germline mutation rate in eukaryotes: a phylogenetic metaanalysis. Evolution Letters, 7(4):216–226, 06 2023. ISSN 2056-3744. doi:10.1093/evlett/qrad027. URL https://doi.org/10.1093/evlett/qrad027.

J. Xu and C. Ané. Identifiability of local and global features of phylogenetic networks from average distances. Journal of Mathematical Biology, 86(1):12, 2023. doi:10.1007/s00285-022-01847-8.

Y. Yu and L. Nakhleh. A maximum pseudo-likelihood approach for phylogenetic networks. BMC Genomics, 16(10): S10, 2015. doi:10.1186/1471-2164-16-S10-S10.

Y. Yu, J. Dong, K. J. Liu, and L. Nakhleh. Maximum likelihood inference of reticulate evolutionary histories. Proceedings of the National Academy of Sciences, 111(46):16448–16453, 2014. doi:10.1073/pnas.1407950111.

G. Zecca, M. Labra, and F. Grassi. Untangling the evolution of american wild grapes: Admixed species and how to find them. Frontiers in Plant Science, 10:1814, 2020.

X. Zhao, Y. Guo, L. Kang, C. Yin, A. Bi, D. Xu, Z. Zhang, J. Zhang, X. Yang, J. Xu, et al. Population genomics unravels the holocene history of bread wheat and its relatives. Nature Plants, 9(3):403–419, 2023.

